# Expression-linked promoter selection (ELiPS) engineers short, strong ubiquitous promoters for gene therapy applications

**DOI:** 10.64898/2026.06.25.734611

**Authors:** Sydney V. Oraskovich, Kazuomori K. Lewis, Joost van Haasteren, Hyuncheol Lee, Esther Chu, David V. Schaffer

## Abstract

Adeno-associated virus (AAV)-based gene therapy has made steady progress towards efficient delivery to numerous target cell populations, yet the virus’s 5 kb packaging limit remains a challenge for effective and in some cases cell-selective cargo expression. Here, we introduce Expression-Linked Promoter Selection (ELiPS), a high-throughput platform for generating and functionally screening >10^6^ engineered, short promoter variants using an AAV expression platform. ELiPS relies on a Golden Gate cloning method to build random oligomers of selected transcription factor binding sites (TFBSs) upstream of a minimal promoter, GFP, and a unique 3’ barcode. As a proof of concept, to engineer short (∼250 bp), synthetic, ubiquitous promoters, we applied ELiPS to build two libraries composed of TFBSs for ubiquitously expressed transcription factors (TFs) and screened them via AAV-mediated transduction *in vitro*. This strategy identified promoters with expression surpassing human cytomegalovirus (CMV) and CAG *in vitro*, and one variant was capable of driving therapeutic expression of B-domain-deleted Factor VIII (BDDFVIII) *in vivo* at levels comparable to a liver-specific promoter benchmark. ELiPS thus establishes a scalable framework for promoter discovery, enabling the design of compact, ubiquitous or cell-selective expression cassettes that enable further precision and efficacy in AAV-based gene therapies.

## Introduction

Gene therapy—the introduction of genetic material into patients for therapeutic benefit—has steadily gained traction as an extremely promising modality to treat human disease. Of the viral and non-viral delivery vehicles employed to date, AAV is among the most effective and potentially safest available^1,2^, as evidenced by numerous AAV-based gene therapy FDA approvals in the last decade. Despite its favorable qualities—such as the parental virus’s non-pathogenic nature, low immunogenicity with local delivery, and long-term transgene expression of over a decade in ongoing studies^3^—the virus faces the shortcoming of delivery of large cargoes such as genome editing tools or cDNAs encoding for example Factor VIII, cystic fibrosis transmembrane conductance regulator (CFTR), dystrophin, and other large proteins^4^. Specifically, genetic cargos cannot exceed ∼5 kb^5^, which for some cDNAs does not enable use of typical ubiquitous, strong promoters (e.g. CMV or CAG). Moreover, in situations where cell selective gene expression is needed to reduce toxicity and/or immunogenicity^6^ and a highly targeted capsid is not yet available^7^, it would be attractive to use a cell-selective promoter, yet their often large size and/or weak expression pose challenges.

Promoter engineering—whether through identifying and harnessing natural enhancers or promoters or through construction of synthetic promoters from known functional motifs^8^—is being actively pursued to identify shorter, stronger, and potentially more cell-specific promoters for a broad range of applications from gene therapy^9^ to synthetic biology. Promoters generally have two major components: a core that contains the elements necessary to recruit RNA polymerase II and one or more enhancers that function as *cis*-regulatory modules (CRMs) to recruit transcription factors that modulate the assembly and activity of transcriptional machinery near the core promoter^10^. As core promoters contain the minimal units necessary to drive transcription, their size is highly conserved across species at approximately 100 bp^11^, and efforts to further reduce this length have not been successful. However, combining elements across multiple species has led to the creation of so-called “super core promoters” (SCPs) that have substantially increased core promoter transcription efficiency^12^. In parallel, enhancer regions modulate promoter function to orchestrate complex cellular processes such as cell division, differentiation, metabolism, and many others^13,14^, and the binding of TFs individually^15^ or synergistically^16^ within a given enhancer strongly influences enhancer function.

Many efforts at promoter engineering involve using next-generation sequencing-enabled approaches to identify natural enhancers with desirable properties and engineer them into artificial promoters. The advent of massively parallel reporter assays (MPRAs) has enabled high-throughput functional profiling of thousands to millions of candidate *cis-*regulatory sequences in mammalian cells through barcode-linked sequencing readouts of reporter expression^17–19^. These approaches have also been applied to dissect regulatory grammar, such as the relationships between transcription factor binding site composition, orientation, and spacing and promoter strength^20–22^. Recent studies have extended MPRA frameworks to synthetic promoter libraries assembled from individual TFBS building blocks, demonstrating the utility of high-throughput combinatorial design coupled with sequencing-based functional readouts^21,23–25^.

One desirable profile is a short, ubiquitous promoter, since the strongest promoters typically used for gene therapy are relatively large, such as the ∼800 bp CMV promoter or ∼1700 bp CAG promoter, which combines the CMV enhancer and chicken β-actin (CBA) promoter^9^. Several efforts have been undertaken to identify minimal ubiquitous promoters capable of driving high levels of gene expression, often through rational design of known enhancers^26–29^. Most existing MPRA implementations rely on plasmid transfection or lentiviral delivery and thus do not fully address the challenge of screening large libraries of compact promoters in their therapeutically relevant episomal AAV context. Notably, while AAV capsid libraries can be screened at million-variant scale^30^, promoter screening within AAV has remained far more limited: no existing platform has achieved ∼10^6^ variant scale, with prior AAV-based reporter assays typically evaluating at most thousands to, in some cases, hundreds of thousands of regulatory elements, and most focus on enhancers rather than fully assembled promoters^31–34^. Additionally, pool-packaged AAV libraries used in MPRAs can lead to chimeric packaging products^35,36^, creating challenges for evaluating promoter candidates and compromising barcode-promoter linkage. We have developed a high-throughput Expression-Linked Promoter Selection (ELiPS) method to generate and screen millions of 200-400 bp promoters in a single experiment. A pool of TFs was identified based on their expression within a given cell type(s), the corresponding double stranded DNA TFBSs were synthesized, an iterative Golden Gate assembly method was used to build oligomers of TFBSs paired with barcodes, and the resulting library was inserted into AAV to enable expression screens via bulk RNAseq or in the future scRNAseq analysis of barcode expression. In a proof-of-concept to demonstrate the utility of the method, we packaged two ubiquitous promoter libraries into AAV— at ∼10^6^ variants constituting to our knowledge the largest AAV-based promoter library to date—and screened them for expression *in vitro*. Following RNAseq, we identified two promoters that outperformed CAG and CMV *in vitro* despite being a fraction of their size. One of these promoters was also capable of producing equivalent levels of BDDFVIII *in vivo* to a standard liver-specific promoter benchmark. We anticipate that this method can be next employed for the identification of synthetic strong and/or cell-selective mini-promoters.

## Results

### Constructing ubiquitous promoter libraries

As any given cell type has a unique repertoire of transcription factors (TFs) that drive gene expression^37^, our platform, termed Expression-Linked Promoter Selection (ELiPS), enables engineering of diverse ∼120 bp multimers of TFBSs that act as enhancers to the super core promoter 2 (SCP2)^12^ (**Fig. 1a**). Interestingly, SCP2 drives little to no activity in the absence of an enhancer, enabling selection of promoters whose TFBS arrays drive expression. Because the strongest and historically most common promoters utilized in gene therapy applications are ubiquitous, as an initial proof of concept demonstration of the utility of this approach we designed two promoter libraries intended to express strongly across a variety of cell and tissue types. First, we selected broadly expressed transcription factors using bioinformatic analysis of the Human Protein Atlas, in which the protein levels of TFs have been quantified across a host of human tissues, down to rare populations of single-cell types^38^. While this analysis allows for the selection of TFBSs based on the presence of the TF proteins themselves, it is not explicitly tied to their likely transcriptional influence. Thus, in a second, complementary approach, selection of TFBSs was accomplished through analysis of the FANTOM5 SSTAR database^39^ in which transcript levels are reported. The associated TFBS-containing regions were then ranked by activity level, and individual candidate TFBSs were chosen. We selected 15 TFBS motifs for Library 1 and 17 TFBSs for Library 2, with several conserved across both libraries, and dsDNA oligonucleotides containing these TFBSs in either forward or reverse orientation flanked by sequences described below for assembly into ELiPS constructs were synthesized (**Sup. Table 1**).

**Figure 1.**
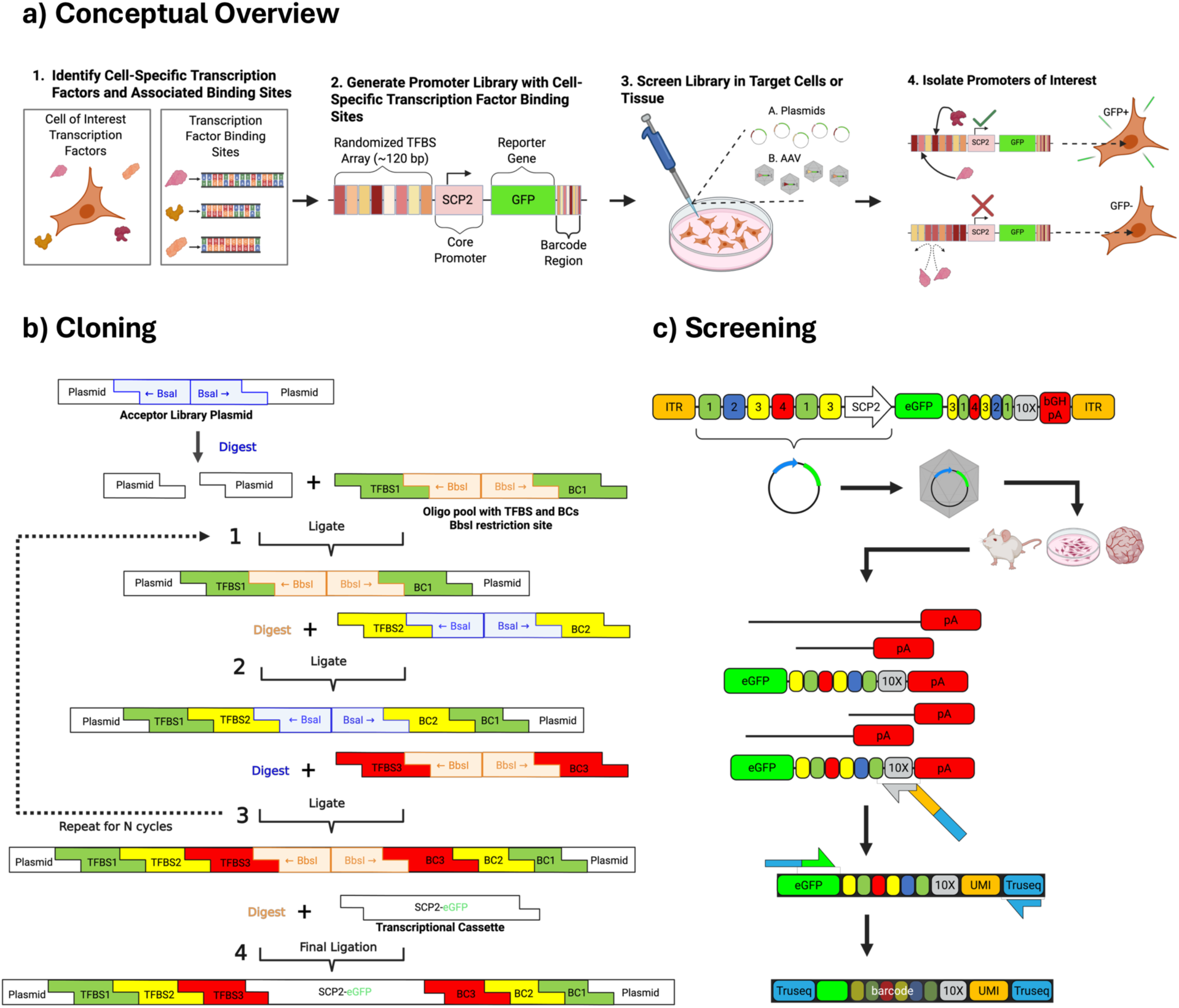
Overview of ELiPS. (a) 1) Transcription factors expressed in the cell or tissue of interest are determined by bioinformatic analysis of characterized TFs from either the Human Protein Atlas (Library 1) or FANTOM5 SSTAR (Library 2) databases. Their binding site sequences are then identified using the JASPAR database. 2) The promoter library is constructed and 3) screened as described. 4) Ideally, a successful promoter is expressed in the desired cell due to a high prevalence of TFs recruited by its enhancer. (b) TFBS sequences in both forward and reverse orientations are synthesized and annealed into dsDNA along with a pair of type II restriction sites (*Bbs*I or *Bsa*I) followed by a unique 4 bp identifier sequence. After pooling the TFBS-containing dsDNAs, iterative Golden Gate cloning is used to build a random multimer where each cloning step inserts a random, new TFBS-identifier pair into the restriction site that lies between the prior TFBS and its identifier. After ∼8 TFBSs and an overall barcode (composed of ten 4 bp identifiers) are assembled via 8 cloning steps, an SCP2-GFP cassette is inserted downstream of the TFBS multimer and upstream of its barcode. Note that dsDNAs with *Bbs*I or *Bsa*I are alternated for each cloning step to eliminate plasmids that did not incorporate a dsDNA oligonucleotide during the prior cloning step. (c) The fully synthesized library, which contains >10^7^ variants of ∼200 bp promoters, is screened in the desired cell or tissue by plasmid transfection or viral transduction. GFP-encoding mRNA is recovered and sequenced to quantify the relative prevalence of each barcode, relative to its prevalence in the original library, and thereby identify the highest expressing promoters.

The ELiPS method assembles a random series or oligomer of these TFBSs upstream of a core promoter, GFP reporter, and a 3’ barcode that can uniquely identify the TFBS oligomer upon NGS analysis of mRNA expression (**Fig. 1b-c**). To construct a library, we performed sequential rounds of Golden Gate cloning to a mixture of the dsDNA oligonucleotides containing the TFBS motifs and a unique 4 bp barcode identifier specific to that TFBS, with an intervening restriction site to enable the next round of Golden Gate cloning^40^. With each cloning cycle into an AAV vector plasmid^41^, one random TFBS/identifier pair is incorporated, thereby exponentially increasing the theoretical library diversity. This approach can thereby emulate some sequence elements of natural enhancers, such as TFBS stoichiometry, order, and orientation, and in the future inter-TFBS spacing as well as heterogeneity in the sequence of individual TFBSs can be varied. For this initial study, we completed N=8 cloning cycles for each library. In the last cloning step, we incorporated an SCP2-GFP cassette, resulting in two ubiquitous promoter libraries with ∼10^6^ clones each.

Prior to screening the full-length library, we first evaluated the ELiPS method using shorter promoter constructs from a smaller pre-defined pool of TFBS/BC oligos. After three TFBS/BC ligation cycles and insertion of an SCP2-GFP cassette, the resulting 3-mer TFBS library was transfected into HEK293T cells (**Sup. Fig. 1a**). Following mRNA extraction and reverse transcription (RT) into cDNA, the barcodes were individually Sanger sequenced to quantify the positional abundance of TFBS motifs alongside barcodes from the plasmid (pre-screened) library. Comparing the positional frequency of each TFBS motif between the pre- and post-screened libraries revealed enrichment of specific TFBS motifs (**Sup. Fig. 1b**). In a subsequent transfection experiment in HEK293Ts, a 3-mer promoter composed positionally of the most enriched TFBS motifs drove stronger levels of GFP than a 3-mer promoter composed of the most-de-enriched motifs and was lower yet comparable in strength to the 183 bp F5tg83 promoter commonly employed as a short promoter in size-limited gene therapy applications^42^ (**Sup. Fig. 1c-d**). Thus, this initial proof of concept established the ability to recover barcoded promoter sequences via targeted RT and successfully assess positional enrichment of specific TFBS motifs in functional promoters.

### Full-length library screening for enriched promoters

Both libraries, generated through 8 cycles of TFBS/BC insertion and confirmed by Sanger sequencing to each be approximately 200 bp, were packaged into an A101 AAV capsid^43^, which strongly transduces HEK293 cells. To ensure that at most one promoter element transduces each cell, thereby mitigating the potential for chimerism events in two or more promoter candidates or biased promoter expression from multiple infections per cell^35^, we first optimized the multiplicity of infection (MOI) by infecting HEK293T cells with an A101-CAG-GFP control virus at varying MOIs (**Sup. Fig. 2**). We fit our results to a Poisson distribution and found that at an MOI of 10,000 each cell was on average infected with 0.76 AAV virions and only ∼18% of cells would be infected with 2 or more AAV virions. We therefore chose to transduce HEK293T cells with our packaged promoter library at an MOI of 10,000. After 72 hrs, RNA was harvested and reverse transcribed, and both the pre- and post-screened library barcodes were subjected to next generation sequencing. For each promoter we calculated the enrichment ratio, defined as the ratio of the number of times a barcode appeared in the expressed mRNA vs. in the original plasmid library.

The promoter elements from both libraries exhibited a broad range of enrichment scores (**Fig. 2a-c**), with only a small fraction of promoters with enrichment scores above 2 (∼12% of promoters in Library 1 and ∼14% of promoters in Library 2). We anticipate that the majority of promoters in each library contained TFBS motifs positioned at sub-optimal spacings, creating steric hindrance and/or competition between neighboring TF binding events and thereby leading to limited promoter activity^44^. Interestingly, a higher percentage of promoters in Library 2 were de-enriched, with enrichment scores less than or equal to 0 (∼72%), relative to promoters in Library 1 (∼45%), suggesting that selection of TF candidates by cellular expression (i.e. the Human Protein Atlas based design for Library 1) may be superior to the selection of TFs for their transcriptional activity (i.e. FANTOM5 SSTAR based design for Library 2). Given that few promoters in each library had high enrichment scores and the low likelihood of multiple AAV infections per cell at our chosen MOI, we further inferred that the chances of a nonfunctional promoter becoming transcriptionally biased by a functional promoter due to chimerism was low. We next examined how the composition, orientation, and spacing of TFBSs within each promoter influenced enrichment. To do this, we compared the probability distributions of individual TFBSs—collapsed across forward and reverse orientations—between the pre-screened plasmid library and the top 5000 most enriched promoter variants (**Sup. Fig. 4**). In Library 1, enriched promoters were overrepresented for TFBSs associated with the AP-1 transcriptional network^45^, including ATF4, FOS, and JUN, with log2 enrichment scores ranging from 0.68 to 1.32. In contrast, Library 2 exhibited a broader depletion of TFBSs, with 13 of 17 motifs appearing at lower frequencies in the enriched set relative to the plasmid library. Despite this overall depletion, a subset of TFBSs was strongly enriched, most notably CREB1 (log2 enrichment = 1.61), NRF1 (0.72), and NFE2L2 (0.62).

**Figure 2.**
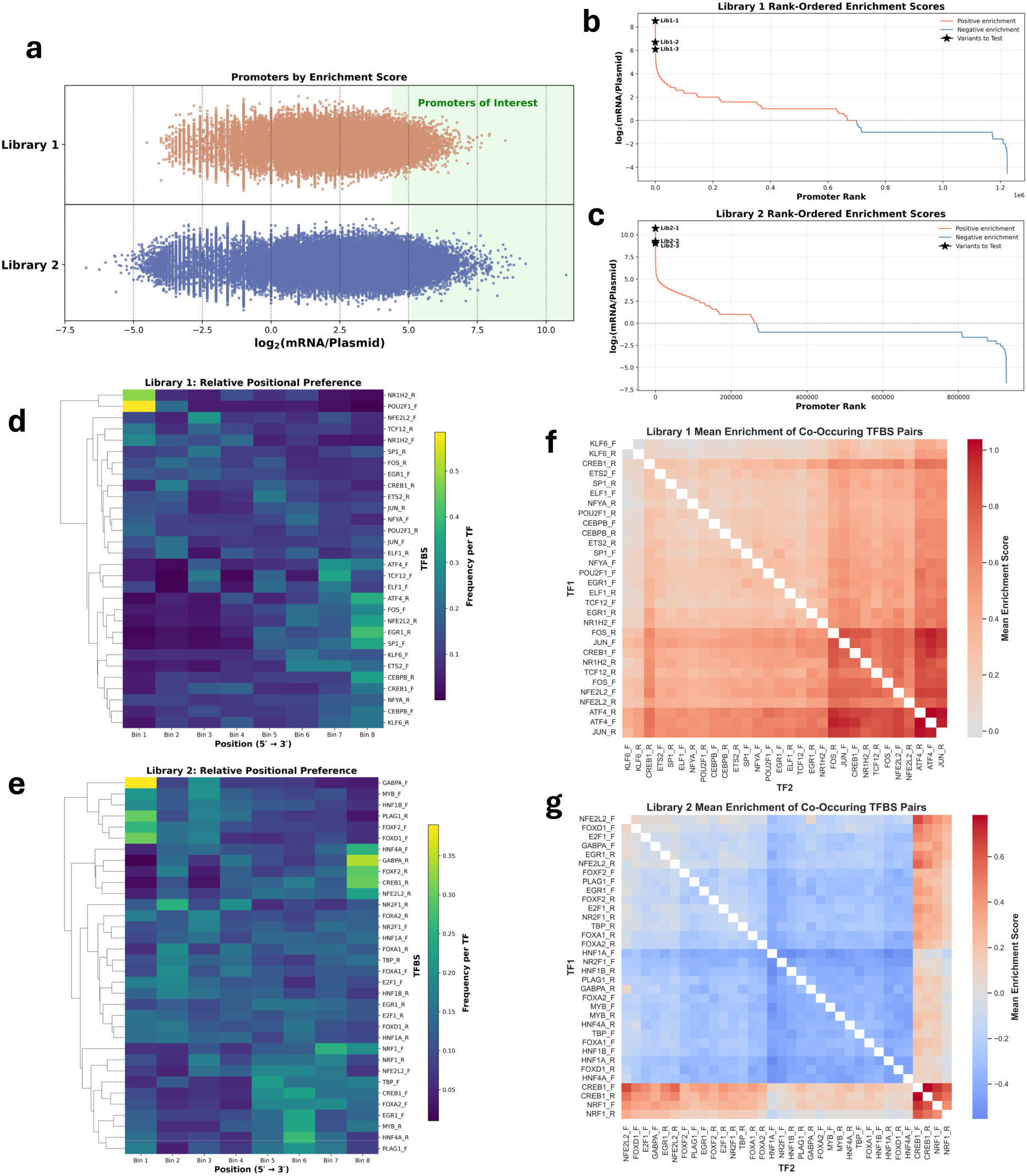
Full-length dual promoter library screen. (a) Following screening of each promoter library in HEK293T cells, pre- and post-screened library barcodes were subjected to next generation sequencing. Library 1 (top) and Library 2 (bottom) promoters are arranged by enrichment score, or the ratio of barcode counts in the expressed mRNA vs. the original plasmid library. The top 5000 promoters from each library are highlighted in green. (b) Promoter rank by enrichment score in Library 1 and (c) Library 2. Positively enriched promoters (red) had barcodes containing higher counts in the post-screened mRNA relative to pre-screened plasmids, whereas negatively enriched promoters (blue) had barcodes containing higher counts in the pre-screened plasmids. Three promoters from each library (starred) were chosen for validation. (d) Heatmap showing the relative positional frequency of each TFBS in the top 5000 most enriched promoters from Library 1 and (e) Library 2. Each TFBS position in the promoter is mapped to one of 8 bins to account for library variation in total number of TFBSs per promoter. The relative abundance of each TFBS in each bin across all promoters is then counted. Rows were normalized per TFBS to reflect positional preference rather than overall abundance. Hierarchical clustering groups TFBSs with similar positional distributions. The positional frequency of each TFBS motif was normalized by its positional frequency in the pre-screened plasmid library. (f) Heatmap depicting the mean enrichment score of promoters containing each pair of TFBSs in Library 1 and (g) Library 2. Each cell represents the average enrichment across all promoters in which the two TFBSs co-occur, independent of spacing or orientation. Hierarchical clustering was applied to reveal TFs with similar combinatorial enrichment profiles.

Differences in TFBS positional frequency—the abundance of any given TFBS at a specific position along the promoter—between all post-screened elements in the top 5000 most-enriched promoters revealed TFBS motifs were differentially enriched after selection (**Fig. 2d-e**). Comparing the positional frequencies with the 5000 most de-enriched promoters from each library (**Sup. Fig. 3**) showed some trends. For example, KLF6_F from Library 1 tended to occur towards the 5’ end of the promoter in the top 5000, but was more prevalent towards the middle of the promoter in the bottom 5000; in Library 2, GABPA_R was highly prevalent at the 3’ end of the multimer in the top 5000 but typically occurred on the opposite end in the bottom 5000. But the positional frequency of each TFBS did not reveal definitive patterns, suggesting that the unique TFBS composition within each promoter—which may confer TF cooperation or steric hindrance depending on the unique TFBS makeup—was more important for promoter function than the preferred position of each individual TFBS.

However, examining the impact of co-occurrence of TFBS pairs on overall enrichment, especially within a positional context, was suggestive of a regulatory grammar contributing to a strong promoter. We first quantified all promoters containing each TFBS pair, irrespective of their relative positions, and calculated the mean enrichment score across promoters harboring each pair (**Fig. 2f-g**). To further assess positional effects, we stratified TFBS pairs based on their spacing within the promoter. Specifically, we identified promoters in which TFBS pairs were immediately adjacent (d=1) or separated by one (d=2) or two (d=3) intervening TFBSs (**Sup. Fig. 5**), and calculated the mean enrichment score for promoters containing each pair at a given spacing and orientation.

In Library 1, pairs composed of bZIP family members^46^—ATF4, JUN, FOS, and CREB1—dominated the top-enriched co-occurrences across all spacing conditions. ATF4 self-pairs (ATF4_F/ATF4_R) and ATF4/JUN co-occurrences were among the highest-scoring pairs at every distance, with mean enrichment scores exceeding 1.0 across d=0 through d=3. Notably, enrichment scores for these bZIP pairs increased with spacing, peaking at d=2 (one intervening TFBS), where ATF4/FOS_R reached a mean enrichment score of 1.26—the highest observed across all conditions in Library 1. This spacing-dependent enhancement suggests an optimal positional context for bZIP cooperativity rather than a simple proximity preference. In contrast, pairs involving KLF6, NFYA, SP1, and ETS2 were consistently among the lowest-enriched co-occurrences across all spacing conditions, with pairs in which two KLF6 motifs co-occurred, or in which KLF6 was paired with NFYA or SP1, reaching strongly negative mean enrichment scores (as low as −0.16 at d=1), indicating suppressive co-occupancy that persisted regardless of spacing.

A comparable pattern was observed in Library 2, where CREB1-containing pairs dominated the high-enrichment co-occurrences across all spacing conditions. CREB1 frequently co-occurred with the oxidative stress-responsive bZIP factors NRF1 and NFE2L2^47^, consistent with their convergent use of CBP as a transcriptional co-activator^48,49^. Enrichment scores for CREB1-family pairs were generally lower at d=0 than at d=1 through d=3, mirroring the spacing-dependent trend observed in Library 1 and further supporting the notion that TFBS proximity alone is insufficient to predict cooperative function. One notable exception was the FOXA1_R/CREB1_F pair, which appeared exclusively as the most enriched combination at d=3 with a score of 0.90, suggesting a spacing-sensitive interaction between forkhead and bZIP factors that manifests only at longer range. Conversely, pairs involving hepatocyte nuclear factors—including HNF1A, HNF4A, NR2F1, and PLAG1—were consistently suppressive, with HNF1A self-pairing and HNF1A/PLAG1 co-occurrence reaching the most negative enrichment scores observed in Library 2 (as low as −0.75 at d=1). Together, these results indicate that promoter enrichment is shaped by both the identity of co-occurring TFBS pairs and their relative spacing, with bZIP-family co-occupancy emerging as a dominant positive regulatory feature across both libraries.

### Testing promoter candidates against benchmark promoters

We selected three highly-enriched promoter candidates from each library for individual validation, including the number one most-enriched promoters from each library, variants from among the top promoter candidates that were well-represented in the pre-screened library (enabling higher statistical confidence), and variants containing enriched co-occurring TFBS pairs including TFBSs from the AP1 complex (ATF4/Jun and ATF4/ATF4 pairs) and CREB1/Nrf pairs (**Sup. Table 2**). We compared our six library hits to CAG (1664 bp), Full-CMV (811 bp), and Mini-CMV (173 bp) using both plasmid transfection and AAV transduction in HEK293T cells. The ELiPS promoters demonstrated high levels of activity: in transfection tests, Lib2-2 was the strongest of the synthetic promoters, performing at 82% of Full-CMV activity and 76% of CAG activity (**Fig. 3a-c**) as determined by flow cytometry. Lib1-3 performed second-best in transfected HEK293Ts, at 71% the activity of Full-CMV and 66% CAG activity. In AAV-based transduction with the A101 capsid, overall expression of the ELiPS promoters was lower relative to CAG and Full-CMV benchmarks, with the best-performer Lib1-2 at 58% the activity of Full-CMV (**Fig. 3d-f**) as quantified by flow cytometry. The synthetic ELiPS promoters were thus able to drive gene expression in transfected HEK293Ts comparable to that of the strongest viral promoters currently used.

**Figure 3.**
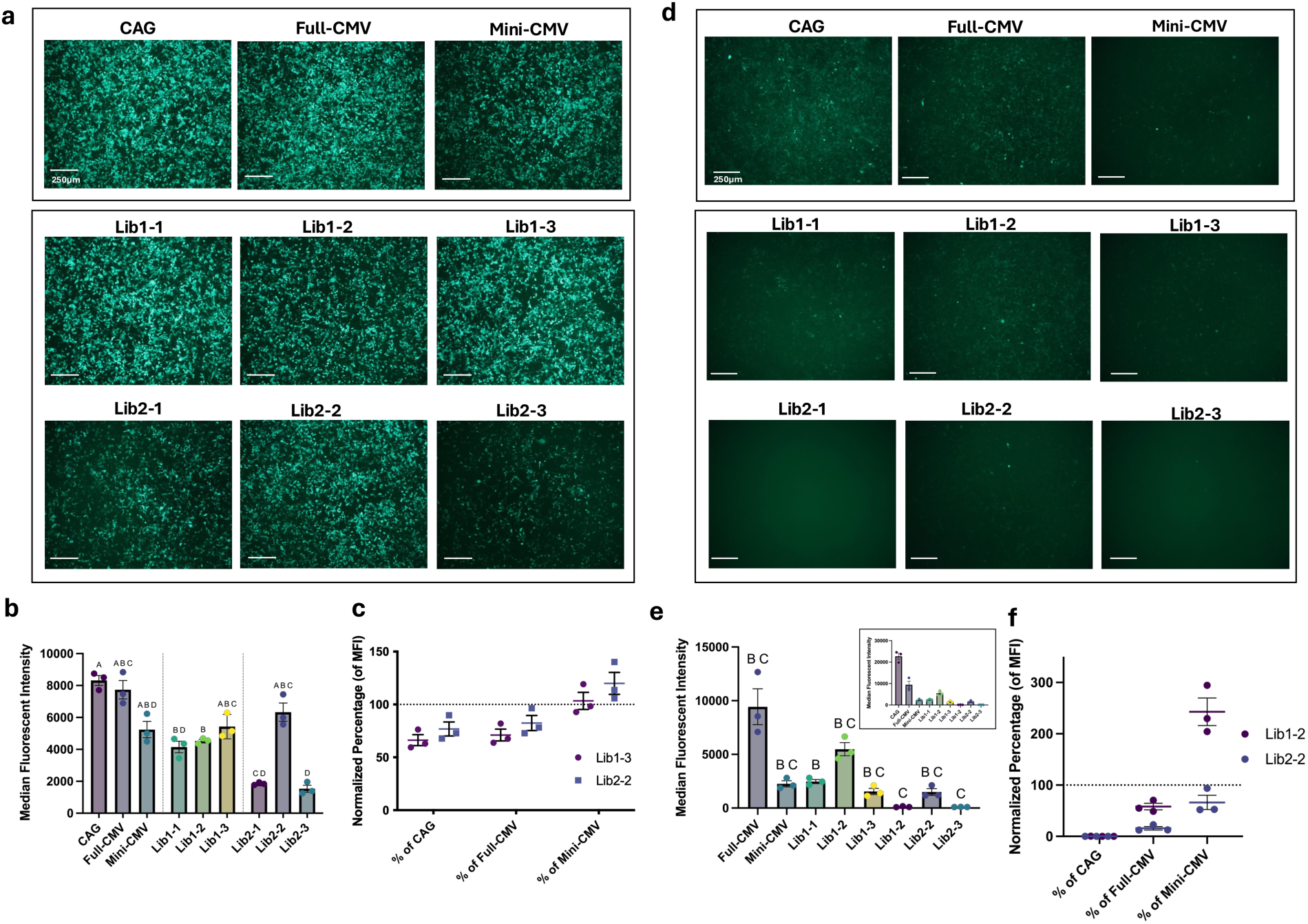
Testing top candidates in HEK293T transfection and AAV-A101 transduction. (a) The three promoters selected for validation from each library were individually cloned and used to transfect HEK293T cells alongside benchmark promoters CAG, Full-CMV, and Mini-CMV. (b) 24 hrs post-transfection, cells were assessed for GFP signal (corresponding to promoter strength) via flow cytometry. (c) Strength of the top performing promoters from each library as a percentage of the benchmark promoters, with gene expression driven by transient plasmid transfection. (d) The same three validation promoters from each library were packaged into an A101 capsid and used to transduce HEK293T cells at an MOI of 20k. (e) 96 hrs post-transduction, cells were assessed for GFP signal via flow cytometry. Inset shows expanded y-axis for low-expression constructs. (f) Strength of the top performing promoters from each library as a percentage of the benchmark promoters, with gene expression driven by AAV transduction. Bars represent mean +/- SEM of per-replicate medians. Statistical differences were assessed using Welch’s one-way ANOVA with Dunnett’s T3 multiple comparisons versus the indicated benchmark. Statistical analyses were performed separately for each benchmark control. Compact letter displays indicate statistical groupings; promoters sharing a letter are not significantly different (adjusted p < 0.05).

### Modular enhancements of ELiPS promoters

Like endogenous mammalian promoters, ELiPS promoters contain an enhancer region upstream of a core promoter, though the enhancer region is considerably shorter (∼120 bp versus hundreds or thousands of bp long). To potentially enhance promoter strength by increasing the number of upstream TFBSs, we thus doubled (designated as −2x) or tripled (designated as −3x) the enhancer region upstream of the core promoter. Additionally, we added intronic elements, which increase transcript stability and mRNA export from the nucleus^50^. In these experiments, we focused on the best performing variants from each library in transduced HEK293Ts: Lib1-2 and Lib2-2. To test the effect of these modular elements on promoter strength (**Fig. 4a**), we compared all Lib1-2 and Lib2-2 constructs against strong ubiquitous viral benchmark promoters through both plasmid transfection and AAV-mediated transduction, as before. In plasmid transfection, the addition of either a double or triple enhancer element significantly increased GFP expression strength by up to 3.3-fold in Lib1-2 and up to 4.7-fold in Lib2-2 (**Fig. 4 b-c** and **Sup. Fig. 6a**). Notably, every tandem enhancer promoter drove expression levels higher than Full-CMV (**Fig. 4d, 4f**) and three of the Lib2-2 promoter variants drove expression levels higher than CAG (**Fig. 4e, 4f**). The largest boost to activity came from the addition of a single extra enhancer unit, and the expression levels of the triple enhancer promoters trended slightly lower, though not statistically significantly different, than those of the double enhancer promoters (p=0.2073 and 0.9403 for Lib1-2 and Lib2-2 double vs triple enhancers, respectively, Welch ANOVA with Dunnett’s T3 post hoc). We speculate this phenomenon may be due to the saturation of local cellular machinery related to the TFBS motifs^51^.

**Figure 4.**
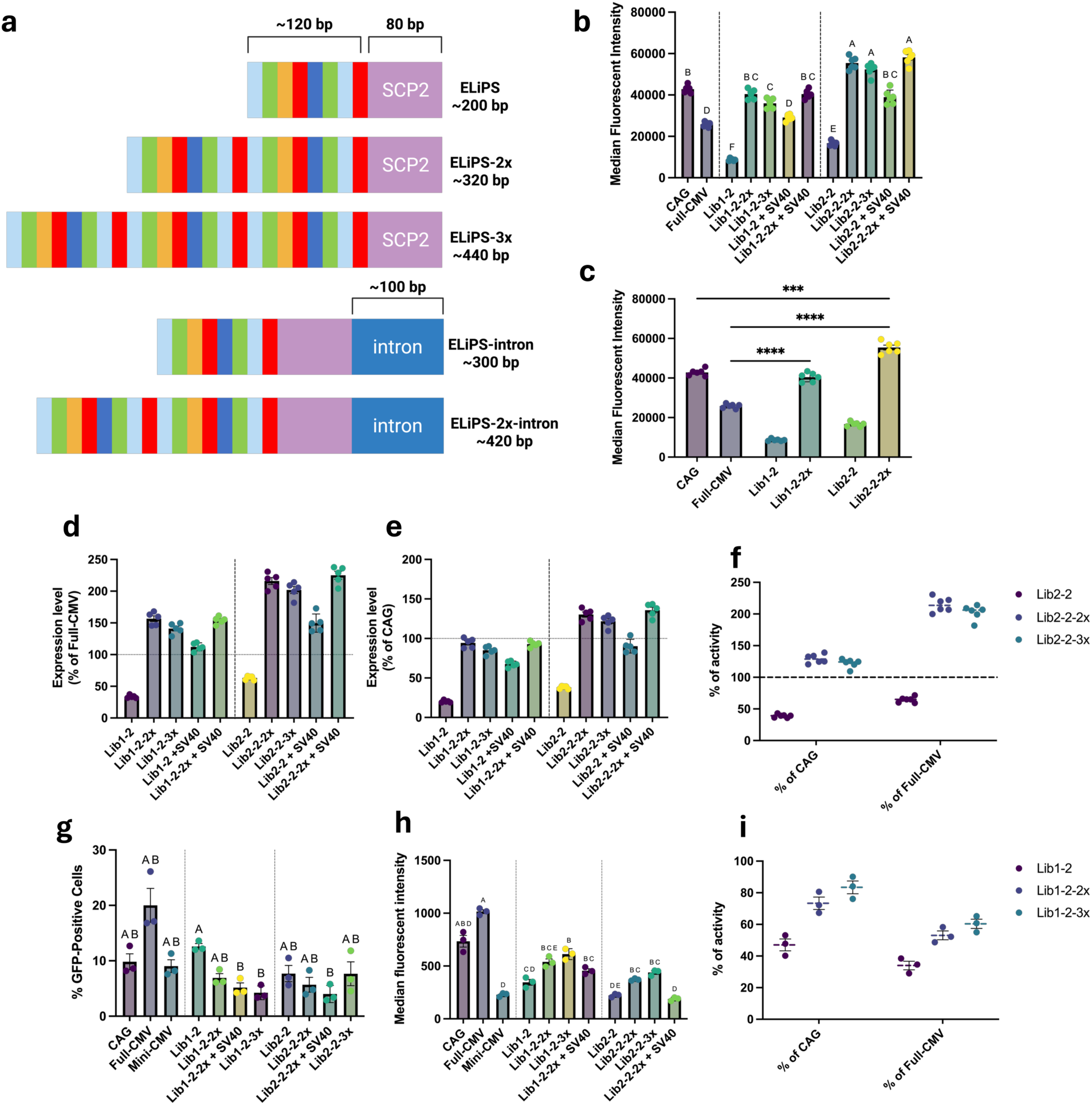
Incorporation of modular elements into top candidates. (a) Visual depiction of the modular enhancer incorporations (2x and 3x) and the intronic elements added to the promoter cassettes. (b) Gene expression driven by transient plasmid transfection of different variants of ELiPS promoters with tandem enhancers and the SV40 intron. (c) Comparison of the base and double enhancer variants of both Lib1-2 and Lib2-2 to controls. (d) The expression levels of promoter variants as a percentage of the strength of the Full-CMV promoter and (e) CAG promoter. HEK293Ts were transfected and analyzed 72 hrs post-transfection. (f) Strength of base and tandem enhancers of best-performing promoter Lib2-2 as a percentage of CAG and CMV activity. (g) Gene expression efficiency measured by the percent of GFP-positive cells driven by AAV- based transduction of HEK293T cells with variants of ELiPS promoters with tandem enhancers and the SV40 intron. Promoters were packaged into the A101 capsid at an MOI of 10,000 and analyzed 72 hrs post-transduction. (h) Promoter strength measured by flow cytometry-based median fluorescent intensity of GFP-positive cells post-transduction. (i) Strength of base and tandem enhancers of best-performing promoter Lib1-2 as a percentage of CAG and CMV activity. Bars represent mean +/- SEM of per-replicate medians. Statistical differences were assessed using Welch’s one-way ANOVA with Dunnett’s T3 multiple comparisons versus the indicated benchmark. Statistical analyses were performed separately for each benchmark control. Compact letter displays indicate statistical groupings; promoters sharing a letter are not significantly different (adjusted p < 0.05). *adjusted p < 0.05 (Welch ANOVA with Dunnett’s T3 post hoc).

The effect of intronic elements was also explored in plasmid transfection. The full-length CMV intron (228 bp), minimal CMV intron^52^ (133 bp), and SV40 intron^53^ (93 bp) were tested. In all cases, both variations of the CMV intron resulted in a decrease of promoter activity (**Sup. Fig. 7**). The inclusion of the minimal CMV intron decreased even the expression of the Mini-CMV promoter, suggesting that truncation of the CMV promoter may remove necessary regulatory elements required for proper use of the intron. However, the SV40 intronic element significantly increased expression levels over the base forms of the promoters by 2-3-fold (**Fig. 4b**). There was, however, only a minimal further increase in expression strength when adding the SV40 intron to the double enhancer promoters. Overall, Lib2-2-2x expressed GFP nearly 2-fold stronger than Full-CMV and, notably, was 30% stronger than the CAG promoter at 25% of its size.

In AAV-based transduction, the additions of enhancer elements and the SV40 intron were also effective at increasing the level of transgene expression over the base formats of the promoters, bringing Lib1-2-3x to 83% the level of CAG expression (**Fig. 4i**). However, in AAV transduction, the Full-CMV promoter has previously been found to be even stronger than CAG across a variety of cell lines^54^, and the ELiPS promoters reached 40-60% of CMV promoter strength following delivery by either the A101 (**Fig. 4g-I** and **Sup. Fig. 6b**) or AAV2 capsid (**Sup. Fig. 8**). Interestingly, the inclusion of the SV40 intron in AAV-based transductions yielded lower expression levels, in one case significantly (p=0.0058 for Lib2-2-2x alone vs the inclusion of SV40, Welch ANOVA with Dunnett’s T3 post hoc), and we surmise that this may be due to differences in splicing and nuclear export of mRNA produced by transfected plasmids vs. episomal viral DNA^55^.

### Testing ELiPS promoters in additional cell lines

The promoter libraries were initially screened in HEK293T cells, and individual validation in this line confirmed high levels of gene expression. However, given the ubiquitous nature of our pre-selected TFs for each library, we hypothesized that the identified promoters may have activity in other cell types. To test this, we transfected a range of mammalian cell lines spanning 3 organisms and 8 tissue models using the double enhancer versions of our top two promoter candidates (Lib1-2-2x and Lib2-2-2x) alongside CAG and Full-CMV benchmarks. The promoters drove strong activity in nearly all cell lines (**Fig. 5** with expression images in **Sup. Fig. 9**) with the exception of SH-SY5Y cells, which are generally difficult to transfect and do not express transcripts from CAG or Full-CMV well. The ELiPS promoters performed especially well in HepG2 cells and CHO-K1 cells relative to benchmarks, with Lib2-2-2x expressing significantly higher than CMV in CHO-K1 and CAG in HepG2 (p=0.0252 and 0.006, respectively, Welch ANOVA with Dunnett’s T3 post hoc). Both ELiPS promoters are thus capable of ubiquitous expression across a variety of cell types.

**Figure 5:**
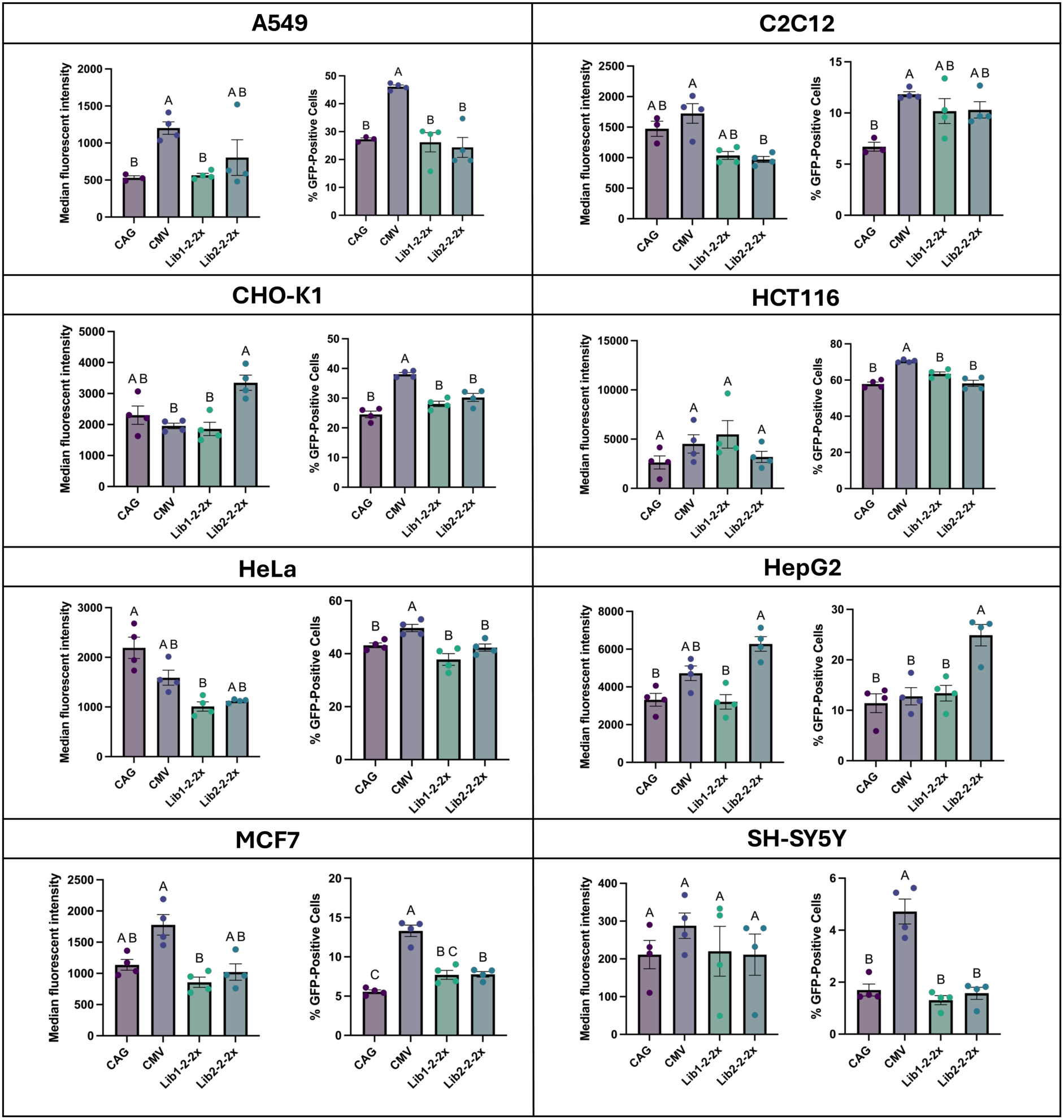
Promoter expression in additional cell lines. Plasmids containing GFP and CAG, Full-CMV, or the double enhancer variants of Lib1-2 and Lib2-2 were each transfected onto a variety of mammalian cell lines. 48 hrs post-transfection, cells were assessed for GFP signal and infectivity rates via flow cytometry. Bars represent mean +/- SEM of per-replicate medians. Compact letter displays indicate statistical groupings; promoters sharing a letter are not significantly different (adjusted p < 0.05 vs benchmark, Welch ANOVA with Dunnett’s T3 post hoc).

### ELiPS promoter-driven BDDFVIII expression *in vivo*

Strong promoter expression *in vitro* does not guarantee utility in animal models, so we were motivated to test our top promoter candidates *in vivo*. We analyzed the capacity of the short promoters to drive expression of a long cDNA that would not fit into an AAV genome with larger conventional promoters such as CAG. One such target is Factor VIII, an essential blood clotting protein whose deficiency is associated with hemophilia A, a genetic bleeding disorder with a global occurrence of nearly 1M individuals^56^. The current standard of care involves multiple weekly intravenous infusions of Factor VIII concentrates–burdensome and costly for patients and their families^57^. While the full-length FVIII coding region spans ∼7 kb, removal of a non-essential B-domain enables protein activity with a size reduction down to 4.4 kb. This B-domain-deleted FVIII construct (BDDFVIII) has enabled packaging and therapeutic expression of the gene into AAV vectors at their maximum packaging limit of around 5-5.2 kb; in fact, this approach became the fourth FDA-approved AAV-based commercial gene therapy (Roctavian by BioMarin)^58^. To enable packaging of BDDFVIII into AAV, the investigators employed a short 252 bp hybrid liver-specific promoter (HLP) known to be highly active in liver cells^59^. Given the high activity of our ELiPS promoters in HepG2 cells, we incorporated our promoters into an AAV-BDDFVIII cassette for comparison to the HLP benchmark.

To first assess how ELiPS promoters fare vs. HLP in a liver derived cell line, we transfected HepG2 cells with CAG, HLP, and our double enhancer variants Lib1-2-2x and Lib2-2-2x. Microscopy and flow cytometry analysis showed comparable expression of the four constructs (**Fig. 6a,b**). We next compared the promoters upon HepG2 transduction with AAV2 (**Fig. 6c**). To enable quantitative analysis in singly infected cells, we transduced at a range of MOIs and fit the data to a Poisson distribution (**Sup. Fig. 10**). While CAG mediated the highest gene expression, we observed significantly higher expression for Lib2-2-2x relative to HLP (p=0.0145, Welch ANOVA with Dunnett’s T3 post hoc). To quantify the capacity to drive protein secretion, we engineered a BDDFVIII-AAV cassette with HLP, Lib1-2-2x, and Lib2-2-2x, incorporating a minimal synthetic polyA sequence previously reported in literature^60^ (**Fig. 6d**). Because wild type FVIII is poorly secreted in the absence of mRNA splicing, we incorporated five amino-acid substitutions (‘X5’) derived from porcine FVIII previously reported to enhance BDDFVIII expression^61^. All Promoter-BDDFVIII-X5 constructs led to ELISA-based detection of BDDFVIII in both transfection and AAV2-based transduction experiments *in vitro* (data not shown), and each construct was thus packaged into AAV8 for liver-aimed murine tropism.

**Figure 6.**
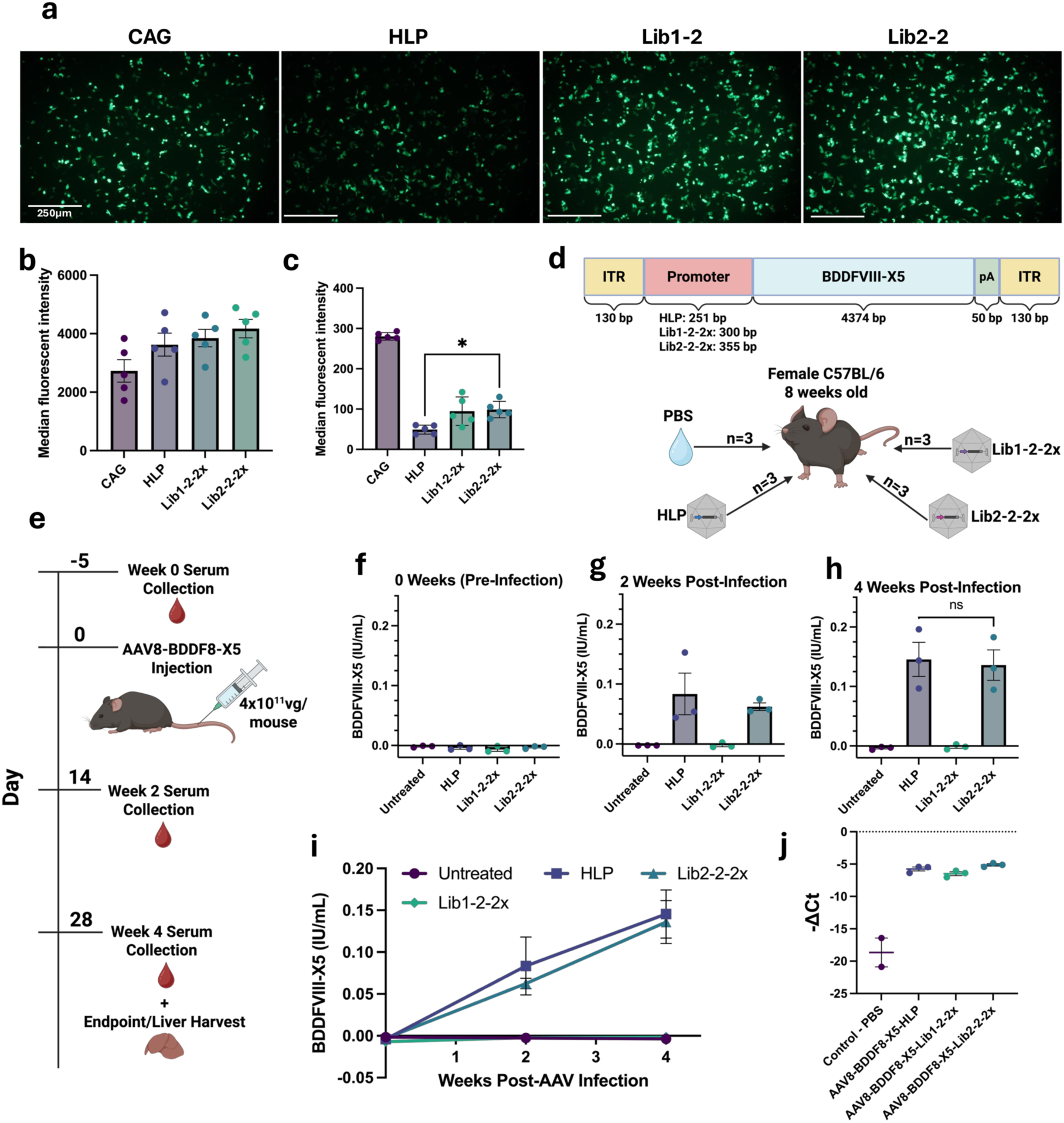
ELiPS promoter-driven expression in a therapeutically relevant context. (a) Visual comparison of GFP-positive HepG2 cells transfected with plasmids containing CAG, a liver-specific benchmark promoter HLP, or ELiPS double enhancer promoters Lib1-2-2x and Lib2-2-2x. (b) Cells were assessed for GFP signal 48 hrs post-transfection via flow cytometry. (c) HepG2 cells were transduced at an MOI of 1000 with AAV2 containing CAG, HLP, Lib1-2-2x, or Lib2-2-2x. 72 hrs post-transduction, cells were assessed for promoter strength via flow cytometry. (d) Representative schematic of the maximized AAV cargo, including two ITRs, the promoter of interest, BDDFVIII-X5, and a minimal synthetic polyA. Each construct was packaged into AAV8 and injected via tail vein into n=3 mice. An additional n=3 mice were injected with PBS as controls. (e) Timeline of the mouse experiment. (f) BDDFVIII levels measured via ELISA from serum samples from each mouse at 0 weeks, (g) 2 weeks, (h) and 4 weeks post-injection. (i) Time course of BDDFVIII serum concentrations throughout experiment. (j) At endpoint, mouse liver samples were harvested and the left lobe of each was isolated and homogenized. Total DNA was extracted from each left lobe and primers specific to human BDDFVIII were utilized in a qPCR-based readout of viral presence, normalized by murine GAPDH housekeeping gene levels. Bars represent mean +/- SEM of per-replicate medians. *adjusted p < 0.05 (Welch ANOVA with Dunnett’s T3 post hoc).

We administered 4×10^11^ vg of each AAV8-Promoter-BDDFVIII-X5 construct (n=3 mice per construct) or PBS via tail vein injection into cohorts of 12 C57BL/6 mice. At 0, 2, and 4 weeks post-infection, mice were bled via submandibular puncture, and serum from each sample was collected (**Fig. 6e**). ELISA revealed that Lib2-2-2x expressed at levels equal to that of HLP (**Fig. 6f-i**). Interestingly, although qPCR quantification revealed that AAV genomes were present in liver at equal levels for each cohort (**Fig. 6j**), Lib1-2-2x-driven BDDFVIII expression was undetectable. In summary, we have demonstrated the ability of Lib2-2-2x to drive expression *in vivo* at levels comparable to a clinically utilized liver promoter, suggesting that the ELiPS method may produce successful promoter candidates for a range of applications.

## Discussion

AAV gene therapy can benefit tremendously from the development of novel classes of synthetic promoters that are small—on the order of 200-400bp—yet retain high levels of activity. The ELiPS platform enables high-throughput MPRA-type screening of such barcoded promoter candidates. This technology addresses the key limitations of AAV-mediated gene therapy applications by enabling the packaging of larger genetic cargos, such as large cDNAs to treat recessive disease or CRISPR/Cas9-based gene editing machinery. This advantage was recently exemplified in a study that used a 50 bp synthetic micro-promoter to accommodate spCas9, a U6 promoter, and a single-guide RNA in a single AAV cassette^62^, however the compact promoter drove diminished expression levels relative to conventional promoters, contributing to a ∼20-30% reduction in editing efficiency. This highlights a central challenge in AAV payload design: size constraints and transcriptional strength are often in tension. The ELiPS platform is uniquely positioned to address both simultaneously, enabling selection of promoters that are not only compact but retain sufficient transcriptional activity. ELiPS also supports the screening of promoter libraries with a diversity ∼10^6^, considerably higher than any existing efforts^31–34^. In this work, we leveraged ELiPS to create two ∼10^6^-element ubiquitous promoter libraries and identified two promoters—Lib1-2-2x and Lib2-2-2x—that can drive expression levels surpassing that of CAG and Full-CMV *in vitro* and express ubiquitously across numerous cell types. We discovered that modular implementation of double enhancer elements significantly boosts expression of top promoter candidates. Finally, we demonstrated the utility of our approach *in vivo* and found that one of our ubiquitous promoters was able to drive comparable AAV-mediated expression of BDDFVIII to a clinically used, liver-selective benchmark. These results establish a promising new tool combining promoter engineering strategies with AAV-based gene therapy optimization.

Despite the considerable advantage of employing MPRAs in AAV engineering and optimization, packaging and screening of large libraries can be confounded by concatemerization of two or more AAV genomes. Furthermore, complex barcoded AAV libraries packaged as a pool can become subject to length- and homology-dependent chimerism due to template switching between adjacent AAV genomes, enabling barcode swapping^35^. Thus, while it is possible that barcode-promoter linkage of our packaged promoter libraries was compromised during packaging and processing, our top-performing promoters selected for validation studies performed well. Additionally, at high coinfection levels, concatemer formation of multiple AAV genomes can permit transcriptional crosstalk between regulatory elements that reside on different genomes^63^. To ensure that AAV genomes remain monomeric episomes, we transduced our libraries at a low MOI, currently the field standard to minimize concatemerization. Combined with the low chance of a strong promoter co-infecting a cell with a weak promoter (based on the enrichment score distribution across each library depicted in **Fig. 2b-c**), transcriptional crosstalk is likely minimal. However, we anticipate that this strategy will be more difficult for AAV-based MPRAs *in vivo*, where MOI is more challenging to control^36^.

Our ELiPS platform also enabled investigation of the regulatory grammar associated with strong or weak promoters—including TFBS composition, orientation, and spacing along the enhancer region. Despite the occasional rare case—such as the IFN-β enhancer—where an enhancer is composed of a highly ordered TFBS arrangement where function is impacted by even slight changes in TFBS spacing^64^, the vast majority of mammalian enhancers are composed of clusters of TFBS motifs with partially flexible spacing and orientation, where motif combinations and density rather than strict positional arrangement encode gene regulatory output^65^. Importantly, our experiments with 2x and 3x tandem enhancers align with past work showing how additional TFBS motifs generally confer a stronger enhancer up to a certain point at which saturation of TF binding occurs^66^. Similarly, we found that our 2x enhancers significantly outperform 1x enhancers, but that our 3x enhancers did not outperform 2x enhancers.

Furthermore, the co-occurrence data from both libraries point to several principles that may reflect underlying features of transcriptional regulatory grammar. First, the consistent enrichment of same-family TFBS pairs—bZIP/AP-1 factors in Library 1 and bZIP/Nrf factors in Library 2—suggests that structural and functional relatedness among co-occurring factors is a reliable predictor of cooperative promoter activity. This is consistent with the well-established tendency of bZIP factors to recognize overlapping or composite DNA elements^67,68^. Second, the observation that mean enrichment scores for bZIP-containing pairs increased from d=0 through d=2 in Library 1 suggests that positional context contributes meaningfully to co-occurrence-based enrichment, and that optimal spacing may exist for particular TFBS combinations. This notion is consistent with literature indicating that both spacing and orientation of binding sites rather than mere presence determines functional output^69,70^. Third, the suppressive pairs identified in both libraries—TFBS co-occurrences associated with strongly negative mean enrichment scores—were notably family-coherent: KLF6 self- and cross-pairs in Library 1 and HNF-family pairs in Library 2 clustered consistently at the bottom of the enrichment distribution. This pattern may reflect competitive occupancy at shared or overlapping binding sites, or alternatively, antagonistic interactions between the pair^71,72^. Finally, the presence of suppressive pairs across all four spacing conditions—in contrast to the spacing sensitivity observed for activating pairs—suggests that the repressive signal associated with certain TFBS co-occurrences may be relatively insensitive to positional context. In future studies, to further understand this regulatory grammar and better design promoters enabling maximal TF cooperation, the incorporation of machine learning models into the ELiPS workflow would be highly desirable. Indeed, recent work has shown that machine learning-guided design of regulatory elements has not only produced stronger elements but uncovered new revelations about transcriptional regulatory grammar^73–76^.

This work has demonstrated the potential of harnessing the ELiPS platform to design mini-promoters for ubiquitous expression, but the technology is flexible for a wider range of applications of TFBS multimers. By drawing on cell-selective activity of numerous transcription factors^77^, ELiPS may be capable of selecting for promoters that can exclusively express in desired cell types. Recent promoter engineering studies have incorporated cell-specific TFBS motifs into regulatory elements to successfully drive cell-selective expression in retinal cells^78^, neurons^79^, and B-cells^80^. Recent advances in single-cell transcriptomics offer new avenues for screening and identifying transcription factor candidates that may contribute strongly to selection of promoters specific to cell identity and/or state, as recently demonstrated in the development of enhancers for hematopoietic progenitor cells^81^. Finally, the ELiPS platform can be expanded outside the realm of gene therapy-centered promoter engineering into any space in which combinatorial arrangement and orientation of regulatory elements drive a desired activity, including CRISPR-based transcriptional regulation^82^, gene circuits for synthetic biology applications^83^, or cell fate engineering and reprogramming^84^.

One unanticipated result from this study was that while two ELiPS promoters, Lib1-2-2x and Lib2-2-2x, drove strong expression *in vitro*, only the latter mediated strong BDDFVIII expression *in vivo*. TF binding and cis-regulatory modules can show substantial conservation across mammalian species^85^, though differences in gene regulation across species^86^ can also impede the translation of promoters designed from human transcriptional datasets to animal models. It may be possible that Lib2-2-2x, which was designed from TFBS motifs associated with TFs that drive strong transcriptional activity broadly across human cell types, may be more translatable to a mouse model due to the conserved and constitutive nature of these TFs. By contrast, Lib1-2-2x was designed from TFBSs whose TFs show enriched expression across human cell types and may therefore be more dependent on cellular context, such as cell-specific signaling or stress states, that may not be equivalently present in the mouse system tested. Alternatively, upon examination of the specific TFBSs in each promoter (**Sup. Table 2**), Lib2-2-2x recruits TFs commonly associated with basal liver function, such as HNF1A and NRF1^87^, whereas Lib1-2-2x recruits TFs previously reported to be associated with some disease states, such as the FOS/JUN AP1 complex and ATF4^88^. The AP1 complex, for instance, expresses at minimal levels in healthy liver yet becomes a master regulator of inflammation in liver disease^89^. This finding prompts us to speculate whether ELiPS promoters can be harnessed to selectively express in diseased cells based on differential TF expression.

In summary, we have developed a high-throughput, AAV-based promoter screening method to generate small, strong promoters. This approach can be further harnessed to investigate how TFBS identities and arrangements contribute to promoter strength, to design cell-specific or even cell-state specific promoters for a range of applications, or design TFBS motif-based elements for applications beyond promoter engineering.

## Methods

### Selection of ubiquitous TFBS motifs

Ubiquitous Library 1: For the first method of TFBS selection through The Human Protein Atlas database, (https://www.proteinatlas.org/), expression values of genes annotated as “Transcription Factors” were downloaded from all available tissues. To find TFs with high and ubiquitous expression, the average of the normalized expression value per gene was calculated for all tissues, totaling 60 tissue types. To select against TFs expressed at very high levels in a small number of tissues, the median and geometric mean were also calculated and only transcription factors with >5 normalized expression values in all three columns were selected for further analysis. A literature search on the resulting transcription factor list was performed, and genes that were implicated in immune responses or could have negative transcriptional activities through post-translational modification were removed from the final TF pool. The top 15 transcription factors ranked by average expression across all tissues were selected for library generation. TFBS motifs (10-14 bp TF binding sequences) were derived from the JASPAR database^90^ to identify their most updated binding motifs or through a literature search.

Ubiquitous Library 2: For the second method of TFBS selection through the FANTOM5 SSTAR database (https://fantom.gsc.riken.jp/5/sstar/Main_Page), mRNA datasets comprising tissue and cell types of interest were analyzed through Cap Analysis of Gene Expression (CAGE) to select TFBS motifs that were over-represented in the proximal region of promoters active in total RNA pool samples. Three different ‘human reference’ mRNA datasets (Clontech Human Universal Reference Total RNA, pool1; SABiosciences XpressRef Human Universal Total RNA, pool1; Universal RNA - Human Normal Tissues Biochain, pool1) were used. TFBS motifs were rank ordered by lowest p-value across replicates (selected TFBS motifs were p < 0.0001). Subsequently, a literature search was performed to remove hits whose associated TFs were implicated in any repressive or inflammatory activity, as well as those requiring protein complexes of larger than 4 transcription factor subunits to drive downstream gene expression. The top 17 TFBS ranked by p-value were selected for library generation, and the most recent versions of each selected TFBS motif were derived from the JASPAR database (http://jaspar.genereg.net/, version 2020).

### ELiPS library cloning

Selected TFBS motifs were designed into oligo sequences for two separate pools (BbsI and BsaI) containing, from 5’ to 3’, (1) an overhang sequence for ligation, (2) the TFBS motif (in either forward or reverse orientation), (3) a restriction enzyme cut site (either BbsI or BsaI), (4) a 4 bp barcode associated with the TFBS motif, and (5) an overhang sequence for ligation (**Sup. Table 1**). Oligos were ordered and diluted to 100 μM, then phosphorylated and annealed in a single thermocycle reaction with T4 DNA Polynucleotide Kinase (New England Biolabs). The oligo pools of all TFBS motifs (differentiated by BbsI or BsaI cut sites) were then pooled in equimolar amounts. A plasmid backbone based on pAAV2/1 (Addgene #112862 from James Wilson) was modified to remove the native *rep/cap* sequences and incorporate an intronic stuffer derived from the human nebulin gene upstream of BsaI acceptor sites, a 10X capture sequence, and a bGH polyA sequence through Golden Gate Cloning (NEB).

ELiPS libraries were cloned as depicted in **Fig 1**. Specifically, 2 μg of ELiPS backbone plasmid was digested with BsaI-HF-v2 (NEB) in 1x CutSmart buffer (NEB) overnight, then run on a 1% agarose gel (SeaKem GTG) and purified with a PureLink PCR Purification Kit (Thermo Fisher). 1 μg of cut plasmid was then ligated to 0.5 μL of the BbsI oligo pool with T4 DNA ligase (NEB) overnight cycling between 25°C and 16°C for ∼80 cycles of 5 min each, followed by heat inactivation at 65C for 10 min. 1 μL of Plasmid-Safe™ ATP-Dependent DNase (Lucigen) was then added with 1 μL of 10 μM ATP and incubated for 1 hr at 37°C, followed by 10 min of heat inactivation at 65°C. The reaction was purified with the Monarch Nucleic Acid Purification Kit (NEB) and transformed into ElectroMax DH10β (ThermoFisher) as per manufacturer recommendations. An aliquot of cells was plated for diversity estimation via colony counting, with the rest cultured overnight in LB supplemented with 100 μg/μL of Ampicillin (Sigma), shaken for 14 hrs at 37°C at 225 rpm. The following day, the cultures were miniprepped using a Wizard Plus SV MiniPrep kit (Promega) and 4 μg plasmid backbone was digested in 4 parallel reactions with BbsI-HF (NEB) in CutSmart buffer for either 1 hr or overnight at 37°C. The reactions were subsequently gel purified as described, and the cut backbone was incubated with the BsaI oligo pool overnight as described. At each step, colonies from the bacterial plates were miniprepped and sequenced with an oligo (5’-TCTGTGAGATACAGAATCAAAAAGC -3’) to confirm linkage between the TFBS sites and corresponding barcodes. These steps were repeated until 8 total oligo ligation steps were complete, after which the plasmid library pool was digested with both BbsI and BsaI and 1 μg was ligated to 1 μg of a SCP2-GFP cassette with compatible overhangs (amplified with the KAPA HiFi PCR kit, Roche, as per manufacturer recommendations). This functional library pool was once again transformed into ElectroMax DH10B cells and prepared for library packaging into AAV using a PureLink HiPure Plasmid Midiprep kit (ThermoFisher).

### AAV library preparation

HEK293T cells were cultured in DMEM (Corning) supplemented with 10% fetal bovine serum (Invitrogen) and 1% Antibiotic-Antimycotic (GIBCO) at 37°C and 5% CO2. AAV libraries for both ubiquitous ELiPS libraries were individually packaged in HEK293T cells as previously described. To package AAV from these lines, HEK293Ts were seeded at a density of 15.2 million cells per 15 cm dish and grown for 72 hrs. Cells were then triple transfected with PEIMAX (Polysciences) using the standard triple transfection technique with the plasmid prep for each ubiquitous library into the A101 capsid, which was previously selected for strong infection of HEK293Ts in the presence of IVIG^43^. In validation studies, individual promoter constructs were packaged into the A101, AAV2, and AAV8 capsids. 72 hrs later, cells were harvested, and the supernatant was collected. The cell pellet was resuspended in 50 mM Tris and 150 mM NaCl, pH 8.5, before undergoing three freeze/thaw cycles (−80°C–37°C). 1 U/mL Benzonase was added, and lysate was incubated at 37°C for 30 min. The lysate was spun at 500 x g for 2 min, and the supernatant was collected and spun again at 3,700 x g for 20 min at 4°C. Virus in the supernatant was precipitated with 8% v/v PEG and 0.5 mM NaCl. After 24 hrs at 4°C, it was spun at 3000 x g for 20 min and the resulting pellet combined with cell lysate.

The virus in crude lysate form was then purified through ultracentrifugation in an Iodixanol gradient. Briefly, ultracentrifuge tubes (Beckman #342413) were loaded with 1.6 mL of 15% iodixanol in PBS-MK (10X PBS, 10 mM MgCl2 and 25 mM KCl), and underlaid with a long needle (Fisher 14-825-16G) with 700 μL of 25% iodixanol, 600 μL of 40% iodixanol, and 600 μL 60% iodixanol, all in PBS-MK. 1.6 mL of lysate was overlaid onto each gradient, and the tubes were spun at 42,000 rpm for 2 hrs at 18°C. Following ultracentrifugation, tubes were pierced at the 40%/60% interface and the bottom 4/5 of the 40% layer and top 1/5 of the 60% layer were removed. The virus in Iodixanol was then purified by spinning through Amicon Pro 100 kDa 50 mL tubes (Millipore-Sigma) 4x with 15 mL of 1x PBS with Tween-20 (Sigma-Aldrich) at 3000 x g for 20 min at 4°C. After concentration down to ∼ 200 μL, the purified virus was stored at 4°C and quantified via real-time quantitative PCR.

### RT-qPCR-based quantification of viral titers

Real-time quantitative PCR was used to quantify the genome-containing viral titer via SYBR Green (Thermo Fisher), 3 mM MgCl2, 0.2 mM dNTP (Roche), and Jump Start Taq (Thermo Fisher) on a CFX RT PCR machine (Bio-Rad). Plasmid DNA was used from a concentration of 10 ng/μL to 0.0001 ng/μL to generate a standard curve with primers against GFP (5’-ACTACAACAGCCACAACGTCTATATCA-3’ and 5’-GGCGGATCTTGAAGTTCACC-3’). Each sample was run with three technical replicates. The resulting titers were used to determine appropriate volumes of virus for transduction of cells in either the library selections or individual promoter validation studies.

### *In vitro* selection for strong promoters in HEK293T cells

To select for strong promoters in 293T cells, both ubiquitous libraries packaged into A101 were transduced onto three biological replicates of 293T cultures in a 24-well plate format at 80% confluency at an MOI of 10,000. After 72 hrs, the RNA was harvested with an RNeasy kit (Qiagen), quantified, and frozen at −80°C.

To prepare the mRNA for next-generation sequencing, 500 ng was reverse transcribed using the iScript Select cDNA synthesis kit (Bio-Rad) along with 500 nM of an equimolar RT oligo pool designed against the 10X capture sequence that contained unique molecular identifiers (UMIs) for demultiplexing PCR duplicates, and the Illumina TruSeq 1 sequence (5 oligos differentiated by the number of N’s to increase diversity for NGS, the N in the following oligo denotes 12-16 total repeats of N; 5’-CTACACGACGCTCTTCCGATCT-N-TTGCTAGGACCGGCCTTAAAGC- 3’). The reaction was then treated with 1 μL of ExoI, incubated for 30 min at 37°C, followed by 5 min at 85°C, then purified with the NEB Monarch kit. This was followed by a phasing PCR with the KAPA HiFi HotStart enzyme and 10 μM of another equimolar pool of oligos that contained homology to GFP sequence, the Illumina TruSeq 2 sequence, and another set of N’s for phasing (N here denotes 0-4 repeats of N; 5’-GTGACTGGAGTTCAGACGTGTGCTCTTCCGATCT-N- GGCATGGACGAGCTGTACAA-3’) for 25 cycles at an annealing temperature of 64.5°C and 15 seconds of extension. Half of the reaction was run on a 1.5% agarose gel and extracted and purified with the Monarch DNA Gel Extraction Kit (NEB). This was followed by Illumina index primer amplification (as per manufacturer design recommendations) with the KAPA polymerase for another 25 cycles at an annealing temperature of 64°C for 5 cycles, then 72°C for 5 cycles. The PCR product was again run on a 1.5% agarose gel and extracted with the Monarch kit. Lastly, the extracted DNA was cleaned up with AmpureXP beads (Beckman Coulter) at a ratio of 1:1.8 for DNA:beads, with 2 washes of 70% EtOH. After quantification via a Qubit Fluorometer (ThermoFisher), samples were submitted for sequencing at the QB3 Vincent J. Coates Genomics Sequencing Laboratory at UC Berkeley. Samples were sequenced with the paired-end 150 bp MiSeq v2 Nano chemistry (Illumina) on a MiSeq machine (Illumina).

### Reconstruction of TFBS motifs and sequences from sequenced barcodes

To convert raw .fastq formatted files from the Illumina sequencing runs into functional TFBS promoter sequences and their associated enrichment ratios, custom python scripts were utilized (available upon request). First, the 150 bp Read 1 and Read 2 sequences from each sample were aligned and merged. Next, PCR duplicates were removed, as designated by any two or more sequences with the same pair of UMIs (12-16 N’s on the 3’ end and 0-4 N’s on the 5’ end). Then barcode sequences from each promoter were extracted from the reads to determine the TFBS motifs and order comprising the promoter that drove expression from each individual mRNA molecule. This output was merged across biological replicates. Enrichment ratios were then calculated by comparing the read count of each individual promoter from the mRNA sample to its prevalence in the plasmid form of each library.

### Cloning of individual promoter fragments

The full-length enhancer sequence identified from the top hits in each screen was synthesized by Integrated DNA Technologies and cloned in the ELiPS backbone in a Golden Gate reaction with an SCP2-GFP cassette. In parallel, the mini-CMV promoter, full-length CMV promoter, and CAG promoter were cloned upstream of GFP. For the tandem enhancer and SV40 intron experiments, 4 or 5-part one step Golden Gate cloning reactions were used to add each element (enhancers x 2-3, intron sequence, SCP2-GFP) into the ELiPS backbone. All constructs were again prepared with the PureLink HiPure Plasmid Midiprep kit (Thermo Fisher). Purified DNA was either used directly for transfection experiments or packaged into AAV as previously described for use in transduction experiments.

### Transfection and transduction in HEK293T cells

To establish transient expression cultures, HEK293T cells were seeded into a 96-well plate at 200,000 cells/cm^2^ and allowed to grow for 18 hrs. They were then transfected with the appropriate promoter construct (5 biological duplicates per promoter) in plasmid format at 500 ng/cm^2^ using PEIMAX (Polysciences) at a DNA:PEIMAX ratio of 1:4 or 1:12. After incubation for 48 hrs, cells were harvested with Trypsin (Corning), washed three times with 1x PBS and assayed on an Attune NxT Flow Cytometer (ThermoFisher) for GFP expression.

For AAV transduction experiments, HEK293T cells were seeded into 96-well format as previously described. After 18 hrs, cells were transduced at a range of MOIs with promoter-GFP constructs in either the A101 capsid (MOI: 10,000–40,000) or AAV2 capsid (MOI: 2,000–5,000). After 72 hrs, cells were harvested with Trypsin (Corning), washed three times with 1x PBS and assayed on an Attune NxT Flow Cytometer (Thermo Fisher) for GFP expression.

All benchmark promoter sequences are depicted in **Sup. Table 3**, and all intron sequences are depicted in **Sup. Table 4**.

### Transfection and transduction in HepG2 cells

HepG2 cells were seeded on a 96-well plate at a density of 3×10^4^ cells/well. 18 hrs after plating, cells were transfected with the appropriate promoter construct using Lipofectamine 3000 (ThermoFisher) at a ratio of 1:4 DNA:Lipo as per the manufacturer’s instructions. For transduction experiments, HepG2 cells were seeded as described and transduced 18 hrs later with the appropriate promoters packaged into AAV2 at MOIs ranging from 50-5000. 48 hrs post-transfection and post-transduction, cells were harvested with Trypsin (Corning), washed three times with 1xPBS and assayed on an Attune NxT Flow Cytometer (ThermoFisher) for GFP expression.

### Flow cytometric analysis

After each flow cytometry experiment, data was processed using FlowJo 10.8 (BD Biosciences). Briefly, cells were gated on FSC H/SSC H for live cells, FSC H/FSC A for single cells, then a BL1 histogram to gate for GFP+ cells. The median fluorescence intensity (MFI) was used to quantify promoter activity of each GFP+ cellular subset, normalized by the MFI of control cell populations. All comparisons were performed using Prism 10 (GraphPad Software). Statistical differences were assessed using Welch’s one-way ANOVA with Dunnett’s T3 multiple comparisons versus the indicated benchmark. Statistical analyses were performed separately for each benchmark control.

### Cloning of the BDDFVIII construct

A FVIII-containing plasmid was procured (Addgene #46774) and cloned via PCR followed by restriction enzyme digest into the previously described AAV backbone using a long 3’ primer that incorporated a 50 bp synthetic polyA sequence previously described in literature^60^ (sequences depicted in **Sup. Table 5**). Promoters of choice were then cloned into the construct. The HLP promoter sequence was isolated from an HLP-containing plasmid (Addgene #109318). To incorporate the X5 amino acid substitutions, a 100 bp oligo containing three of the additions was cloned into the FVIII gene, and a Q5 Site-Directed Mutagenesis Kit (NEB) was used for the remaining two substitutions. The resulting BDDFVIII-X5 constructs were packaged into AAV2 for *in vitro* validation and later into AAV8 for *in vivo* experiments.

### Promoter-driven BDDFVIII expression in mice

A total of 12 C57BL/6 mice were used for the study (JAX), and all mice were female to minimize mouse-to-mouse aggression throughout the study. The 7-week-old mice were given a week to acclimate, then a lancet was used to extract 100-200 μL of blood from each mouse via submandibular puncture. Extracted blood was immediately placed in 3.2% sodium citrate anticoagulant (Fisher Scientific) at a ratio of 1:9 anticoagulant: blood. All extracted blood was then centrifuged at 1500 x g for 15 min, and the serum supernatant was removed and stored at −80°C for the remainder of the study. Blood was collected in the same manner 2 weeks post-infection and 4 weeks post-infection.

Five days after the first bleed, each mouse received a tail vein injection of 100 μL of 4×10^11^ vg AAV8-BDDF8-X5 concentrated in PBS (n=3 mice for each promoter, randomly chosen) or PBS only (n=3 mice, randomly chosen). 4 weeks post-infection, mice were sacrificed and livers were collected and stored in RNA Later Stabilization Solution (ThermoFisher). The left lobe of each liver was isolated, homogenized with a needle, and total DNA was harvested using a DNeasy Blood and Tissue Kit (Qiagen).

A human FVIII-specific primer (5’-GGGAAATCAAGACTCCTTCACACC-3’) and a polyA primer (5’-AAGATCTTTTATTTCAGTAGAGGTCCTGT-3’) were used in a qPCR reaction to quantify viral presence in each liver sample. A second pair for primers specific to the beginning of the human FVIII gene (5’-GAATCTGTAGATCAAAGAGG-3’ and 5’-AAACTATCAAAAACATAGCC-3’) were used to repeat the experiment, and all CT values were normalized by murine GAPDH (primers 5’-AAGGTCGGTGTGAACGGATTTG-3’ and 5’-CCGTGAGTGGAGTCATACTGG-3’).

To measure the concentration of FVIII in each serum sample, the Factor VIII Antigen Plus Kit was used (Affinity Biologicals). Serum was diluted 1:5 in sample diluent, otherwise all steps in the manufacturer’s protocol were followed. Prior to the onset of the mouse experiment, the kit was used to confirm the presence of FVIII in cell media samples following transfection and transduction with each BDDFVIII-X5 promoter construct; here, 100 μL of cell culture media was used for each sample with no dilution in sample diluent.

## Acknowledgements

Library preparation (Qubit quantification and fragment length analysis) and sequencing was done by the Functional Genomics Lab (FGL) and the Vincent J. Coates Genomics Sequencing Lab (GSL) at the University of California, Berkeley. Flow cytometry was performed at the QB3 Cell & Tissue Analysis Facility at the University of California, Berkeley. We thank the UC Berkeley DNA Sequencing Facility for providing Sanger sequencing data and the Cell Culture Facility for providing cell lines for transfection studies. We’d also like to thank Mary West for flow cytometry training and support.

## Author Contributions

DVS supervised the entire project. JvH designed the intial method of cloning ELiPS libraries and optimized it with KKL. JvH and KKL cloned the two ubiquitous libraries and ran the selection in 293T cells. SVO wrote the Python analysis scripts and analyzed the data for the 293T promoters. KKL and SVO performed most validation experiments, analyzed the data, and wrote the manuscript. SVO designed and performed all FVIII experiments and prepared the manuscript for submission. HL performed many of the procedures in the mouse study, and EC assisted throughout the mouse study and endpoint validation.

