## Supplementary Figures for "Expression-linked promoter selection (ELiPS) engineers short, strong ubiquitous promoters for gene therapy applications"

### Supplementary Figure 1

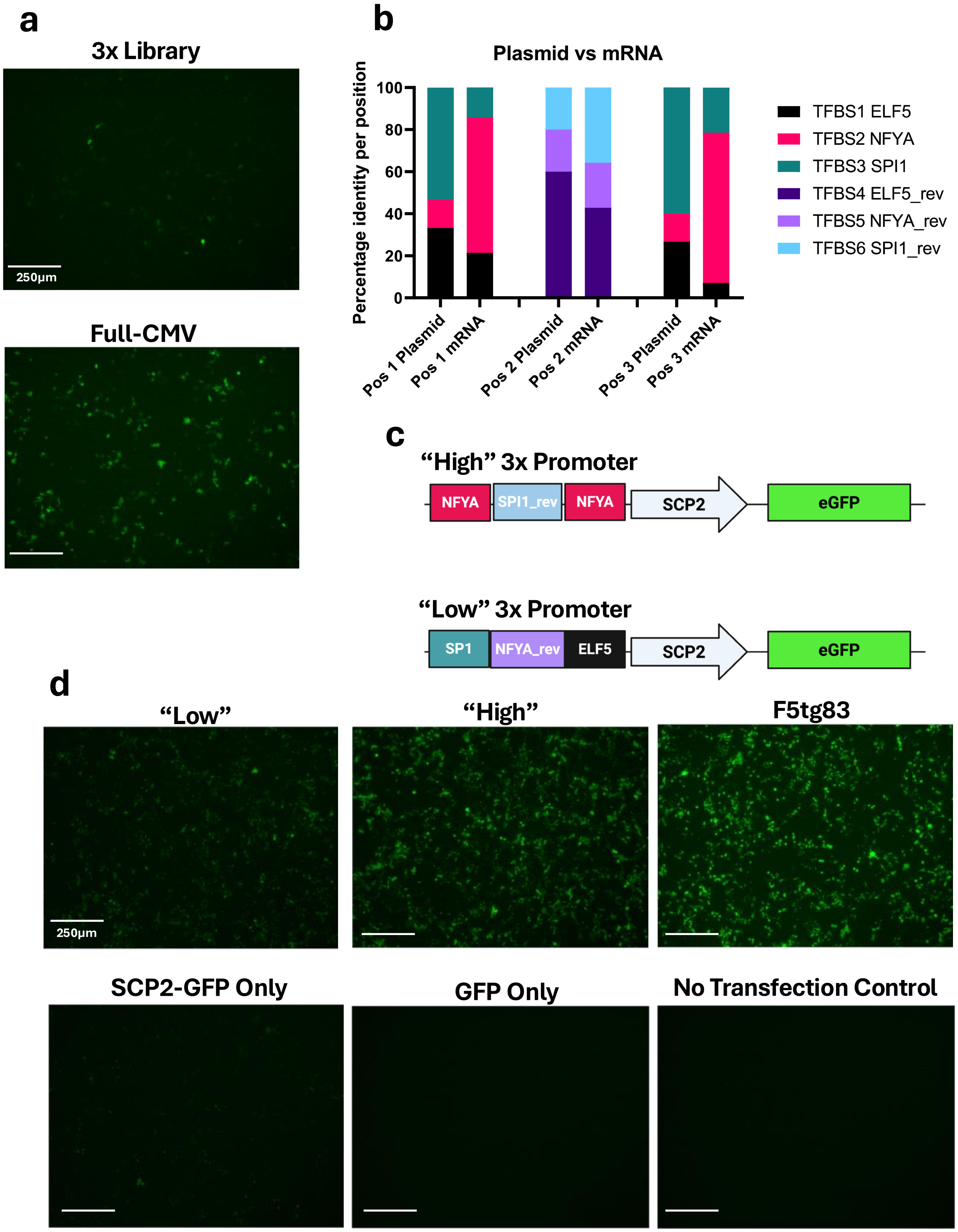

**Supplementary Figure 1. Barcoded promoter screening proof-of-concept.** (a) HEK293T cells were transfected with a minimal 3xTFBS promoter library and GFP expression was visually compared to a Full-CMV-transfected benchmark. GFP signal of the cells transfected with the 3x library was lower than CMV-transfected cells, though there were recognizably bright cells, indicating the ~120 bp promoters were capable of expression at a fraction of the size of the 811 bp Full-CMV promoter. (b) Positional abundance of TFBS motifs in pre- and post-screened promoters. Some TFs, such as NFYA, had relatively low abundances in the pre-screened library and high counts in the sequenced mRNA, indicating motif-specific enrichment. (c) Sequence identity of 'high' and 'low' promoters derived by motif abundance in post-screened promoters. (d) GFP expression of individual clones from the 3xTFBS library. Promoters containing highly abundant post-screened TFBSs exhibited stronger levels of GFP expression in HEK293T cells via transfection.

#### Supplementary Figure 2

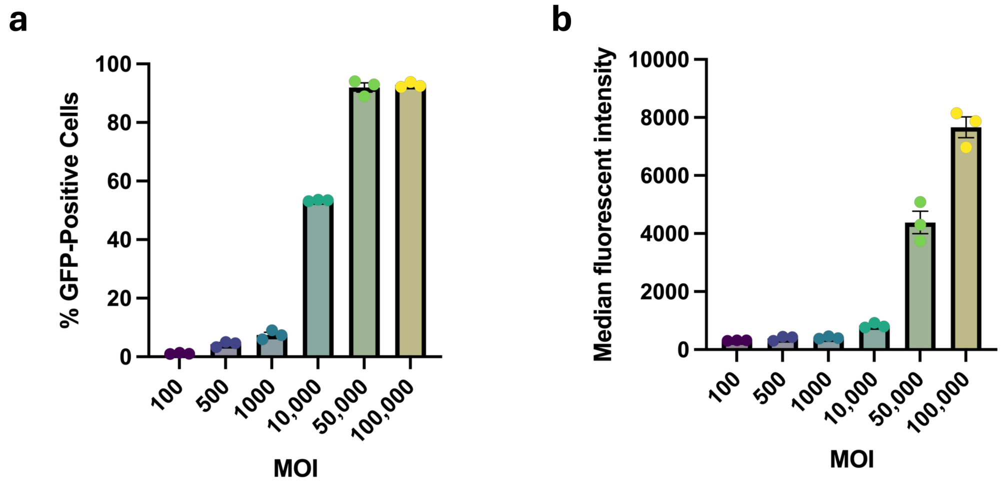


**Supplementary Figure 2. A101-mediated HEK293T CAG-GFP transduction efficiency by MOI.** To ensure the screened promoters would not be subject to recombination due to two or more infection events in a single cell, CAG-GFP constructs were packaged in an A101 capsid and transduced onto HEK293T cells at a range of MOIs. 48 hrs post-transduction, the infectivity rates (a) and GFP expression strength (b) was assessed via flow cytometry. Bars represent mean +/- SEM of per-replicate medians.

### Supplementary Figure 3

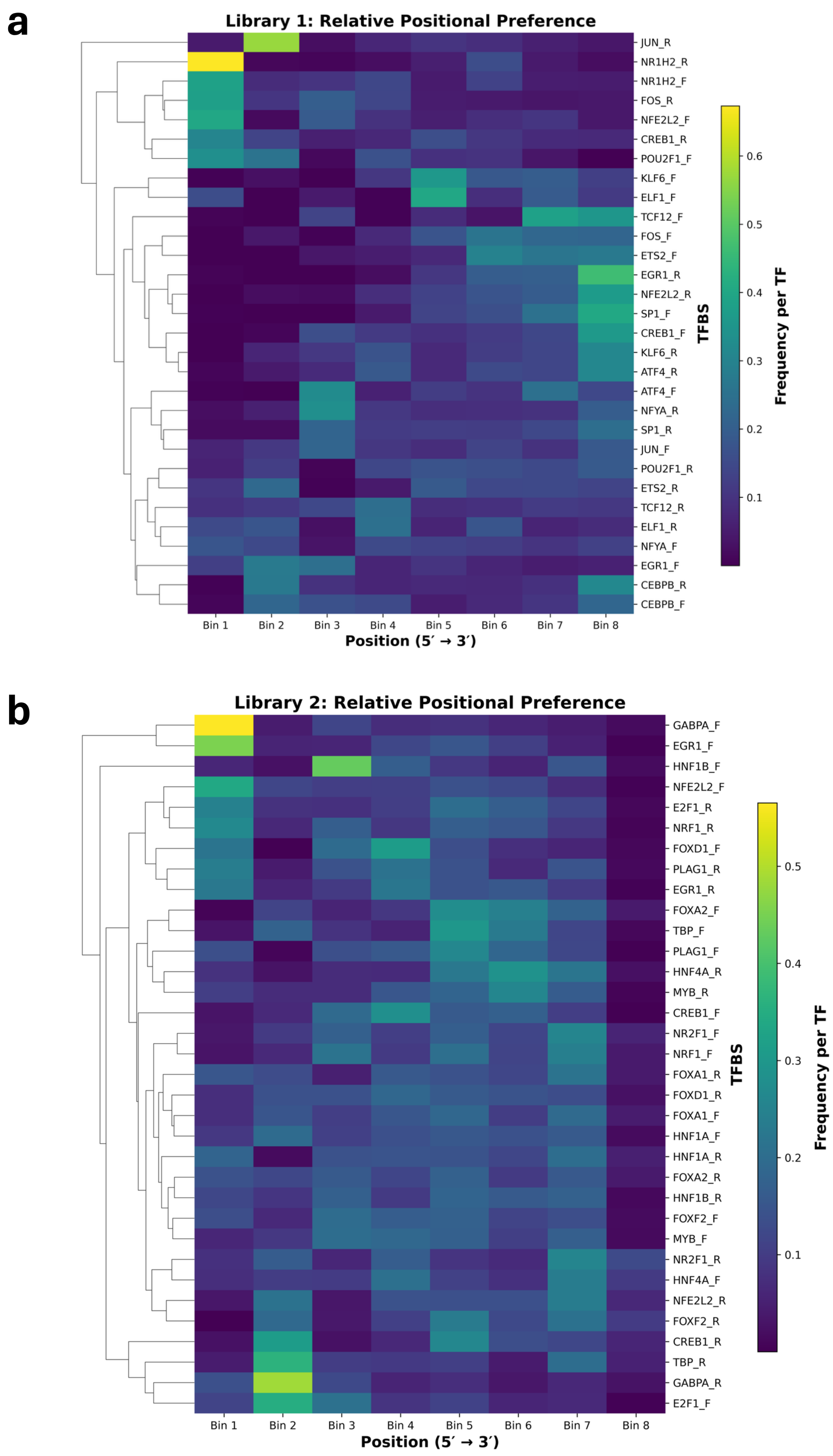

**Supplementary Figure 3. Positional frequency of bottom 5000 promoters in each library.** (a) Heatmap showing the relative positional frequency of each TFBS in the 5000 least enriched promoters from Library 1 and (b) Library 2. Each TFBS position in the promoter is mapped to one of 8 bins to account for library variation in total number of TFBSs per promoter. The relative abundance of each TFBS in each bin across all promoters is then counted. Rows were normalized per TFBS to reflect positional preference rather than overall abundance. Hierarchical clustering groups TFBSs with similar positional distributions. The positional frequency of each TFBS motif was normalized by its positional frequency in the pre-screened plasmid library.

#### Supplementary Figure 4

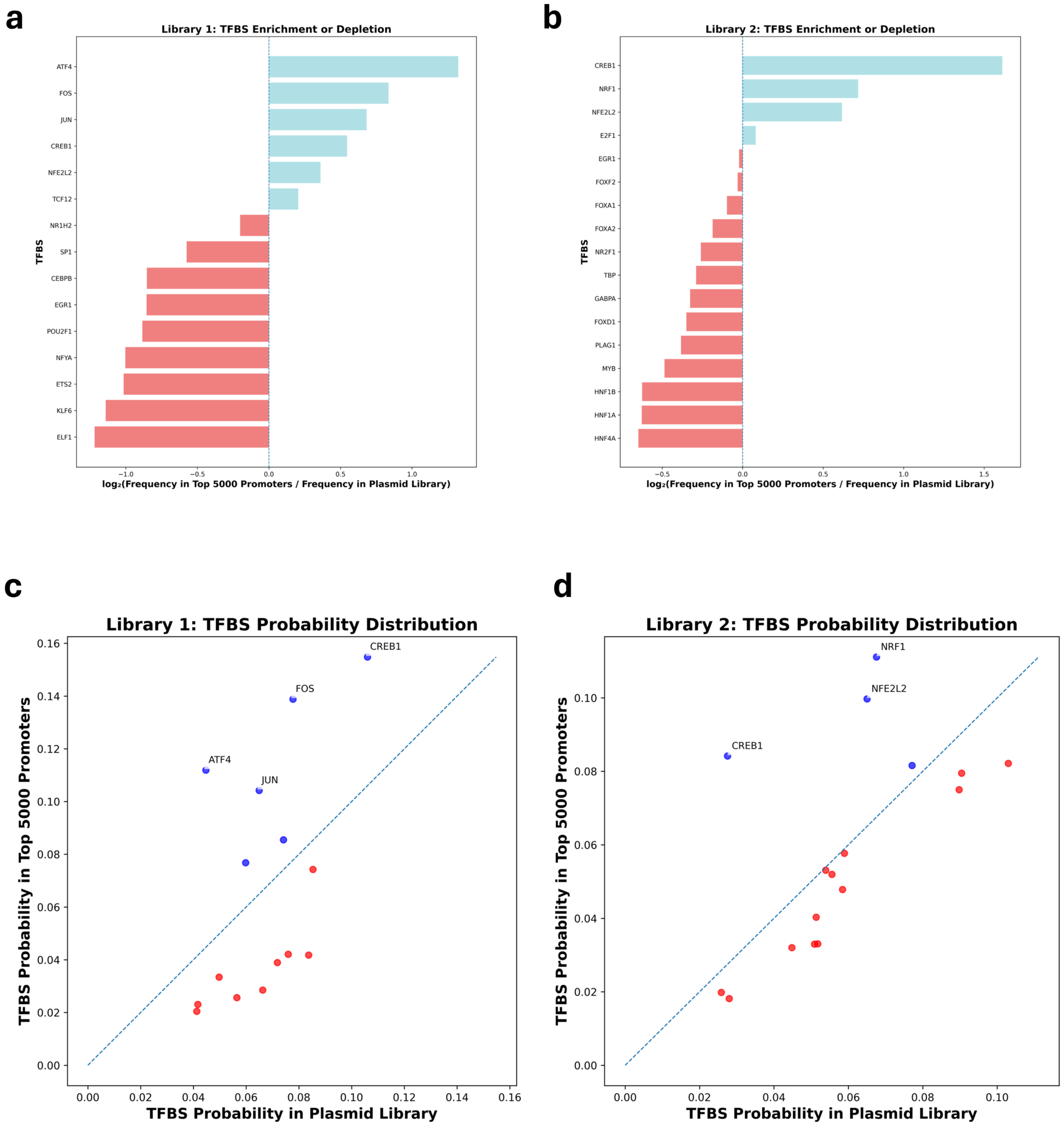


**Supplementary Figure 4. TFBS probability distribution between pre- and post-screened promoter libraries.** (a) Bar plot depicting enriched (blue) and depleted (red) TFBSs in Library 1 and (b) Library 2. The log fold-change in TFBS frequency was calculated by comparing the probability of each TFBS in the top 5000 most enriched promoter variants to its probability in the pre-screened plasmid library. TFBS frequencies were computed based on total motif occurrences across all promoter constructs, weighted by sequence counts. TFBS identifies were collapsed across motif orientation (forward and reverse complements) prior to analysis. (c) Scatter plots showing probability distributions of each TFBS in Library 1 and (d) Library 2, with frequency in the pre-screened plasmid library plotted on the x-axis and frequency in the top 5000 most enriched promoters on the y-axis. The dashed diagonal line indicates equal representation between the two datasets. TFBSs above the diagonal (blue) are enriched, whereas those below the diagonal (red) are depleted. Selected TFBSs of interest are annotated.

### Supplementary Figure 5

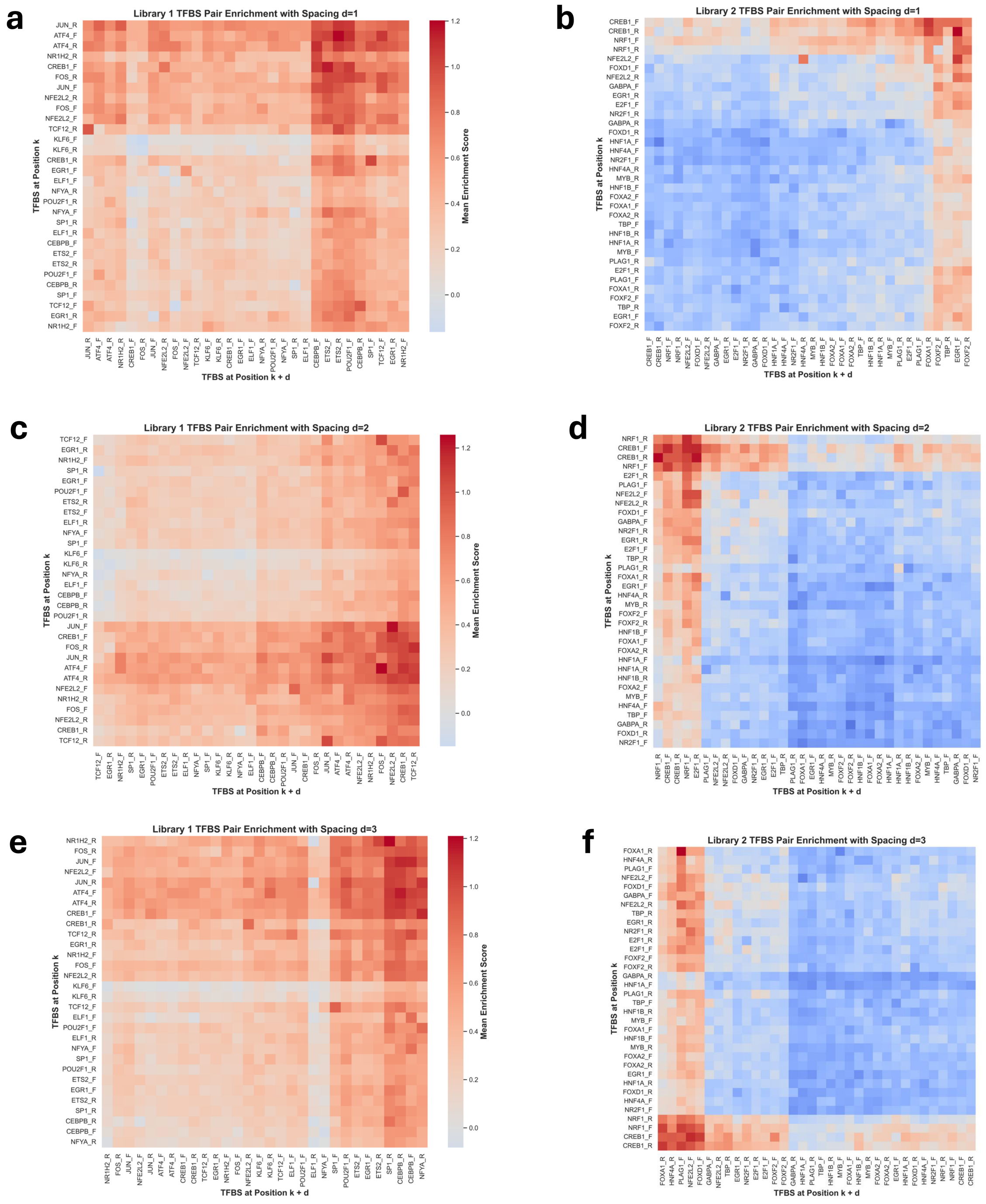


**Supplementary Figure 5. Enrichment by position of TFBS pairs.** (a) Heatmap depicting the mean enrichment score of promoters containing each pair of TFBSs in Library 1 and (b) Library 2 spaced immediately adjacent to each other (d=1), (c, Library 1; d, Library 2) with an additional TFBS between the pair (d=2), or (e, Library 1; f, Library 2) with two additional TFBSs between the pair (d=3). Each cell represents the average enrichment across all promoters in which the two TFBSs co-occur at the designated spacing d=1, 2, or 3, independent of forward or reverse orientation. Hierarchical clustering was applied to reveal TFs with similar combinatorial enrichment profiles.

#### Supplementary Figure 6

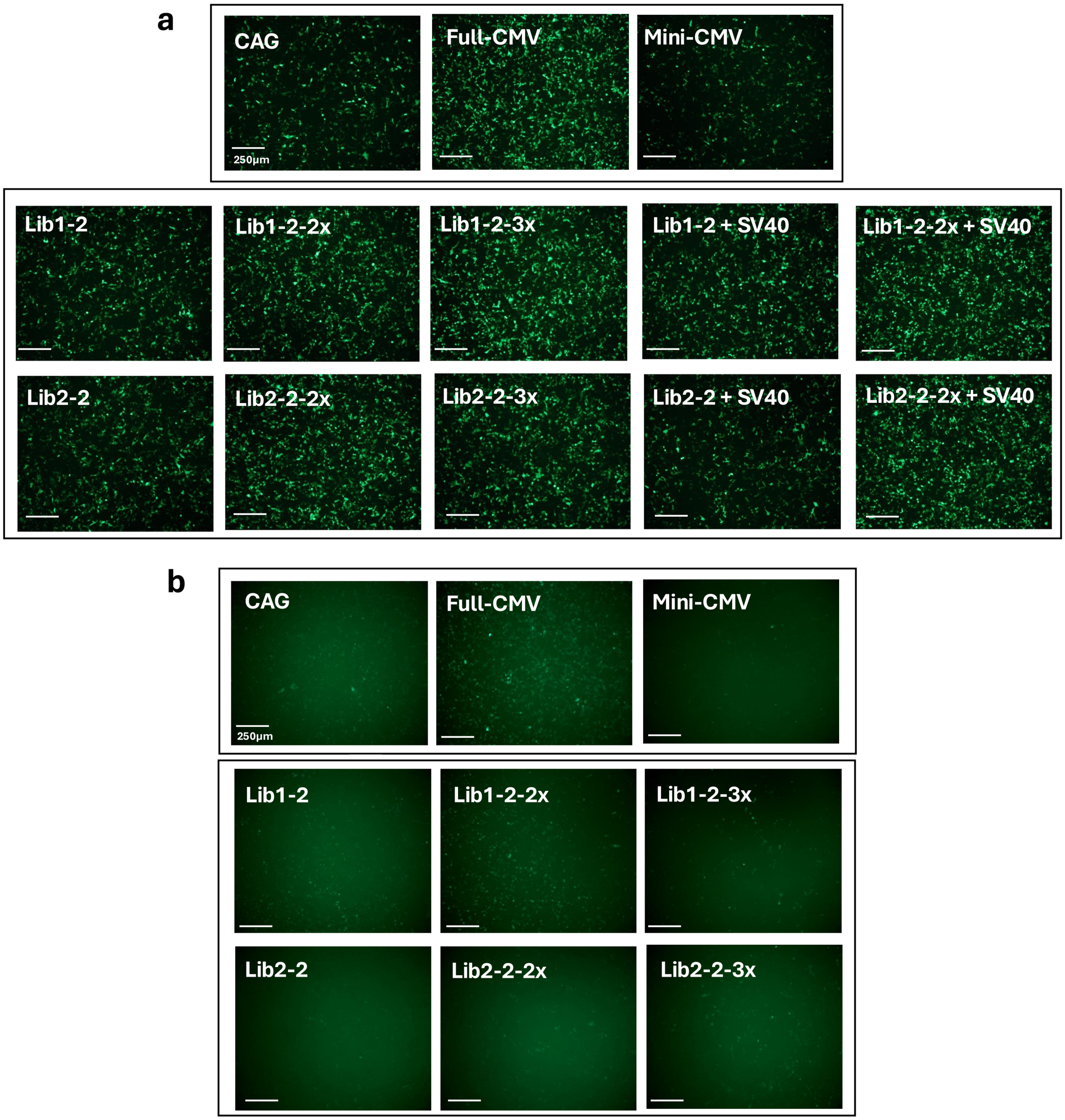


**Supplementary Figure 6. GFP fluorescence with modular elements – transfection and A101-mediated transduction.** (a) Images depicting GFP fluorescence from various promoter constructs captured 48 hrs post-transfection and (b) 72 hrs post-transduction with constructs packaged in A101 and transduced at an MOI of 10,000.

#### Supplementary Figure 7

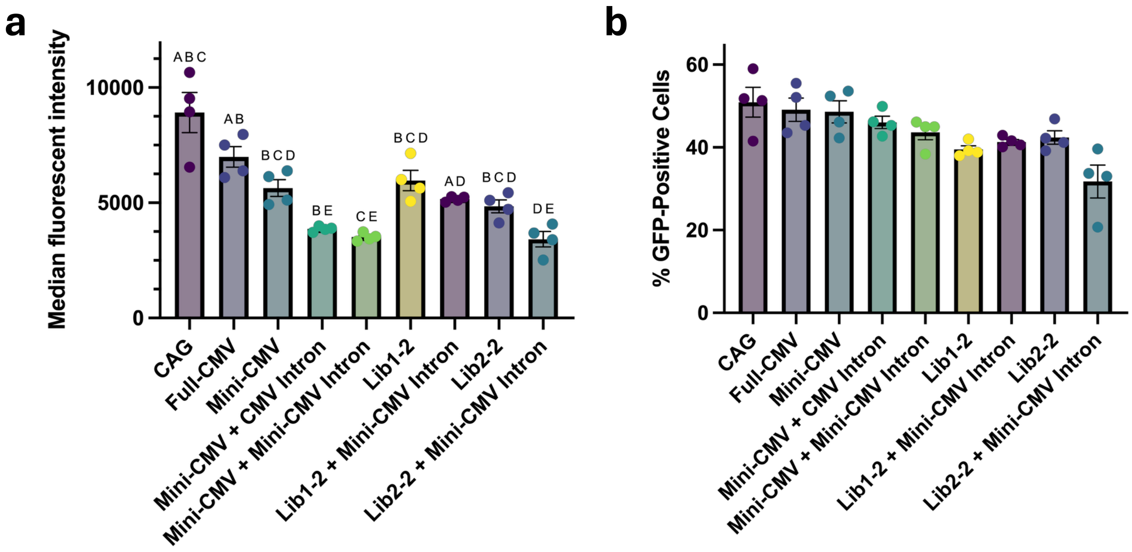


**Supplementary Figure 7. Promoter modularity with Full- and Mini-CMV introns.** (a) Promoter strength of ELiPS promoter containing full-length CMV intron is decreased after transient plasmid transfection of HEK293T cells, despite having relatively similar GFP+ cell percentages (b). Cells were transfected with each construct and analyzed for GFP expression 48 hrs later. Bars represent mean +/- SEM of per-replicate medians. Compact letter displays indicate statistical groupings; promoters sharing a letter are not significantly different (adjusted $p < 0.05$ vs benchmark, Welch ANOVA with Dunnett's T3 post hoc).

#### Supplementary Figure 8

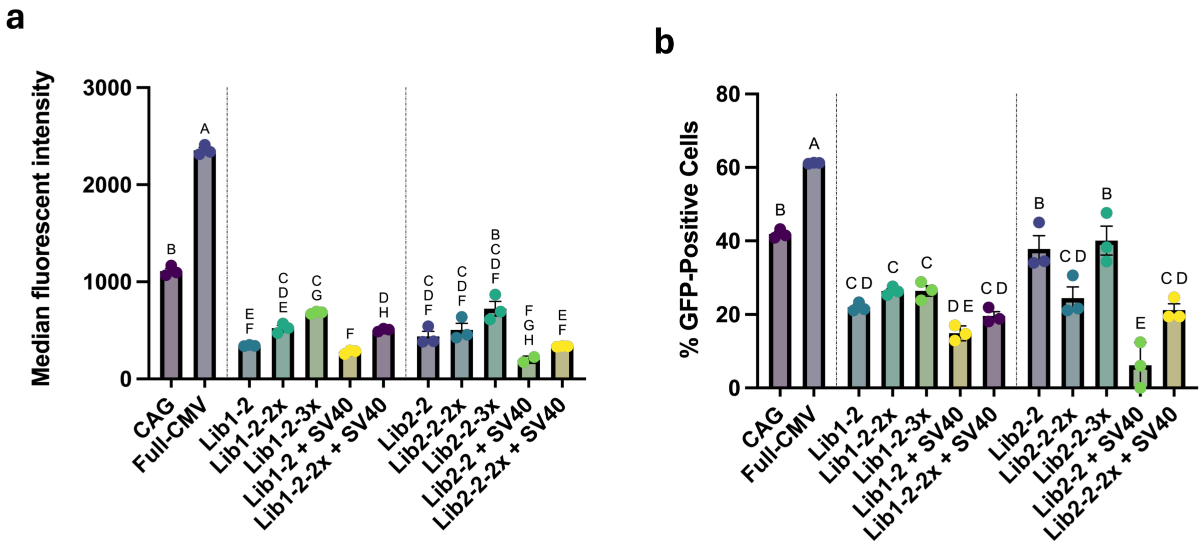


**Supplementary Figure 8. Promoter modulatory with AAV2 transduction.** (a) The overall strength of ELiPS promoters is lower when they are packaged into the AAV2 capsid and used to transduce HEK293T cells. (b) GFP+ cells resulting from transduction with ELiPS promoters and controls. The initial ELiPS promoters were selected with the A101 capsid and have stronger expression profiles when the A101 capsid is used for transduction. Cells were transduced at an MOI of 2,000 with the AAV2 capsid and analyzed for GFP expression 72 hrs later. Bars represent mean +/- SEM of per-replicate medians. Compact letter displays indicate statistical groupings; promoters sharing a letter are not significantly different (adjusted $p < 0.05$ vs benchmark, Welch ANOVA with Dunnett's T3 post hoc).

#### Supplementary Figure 9

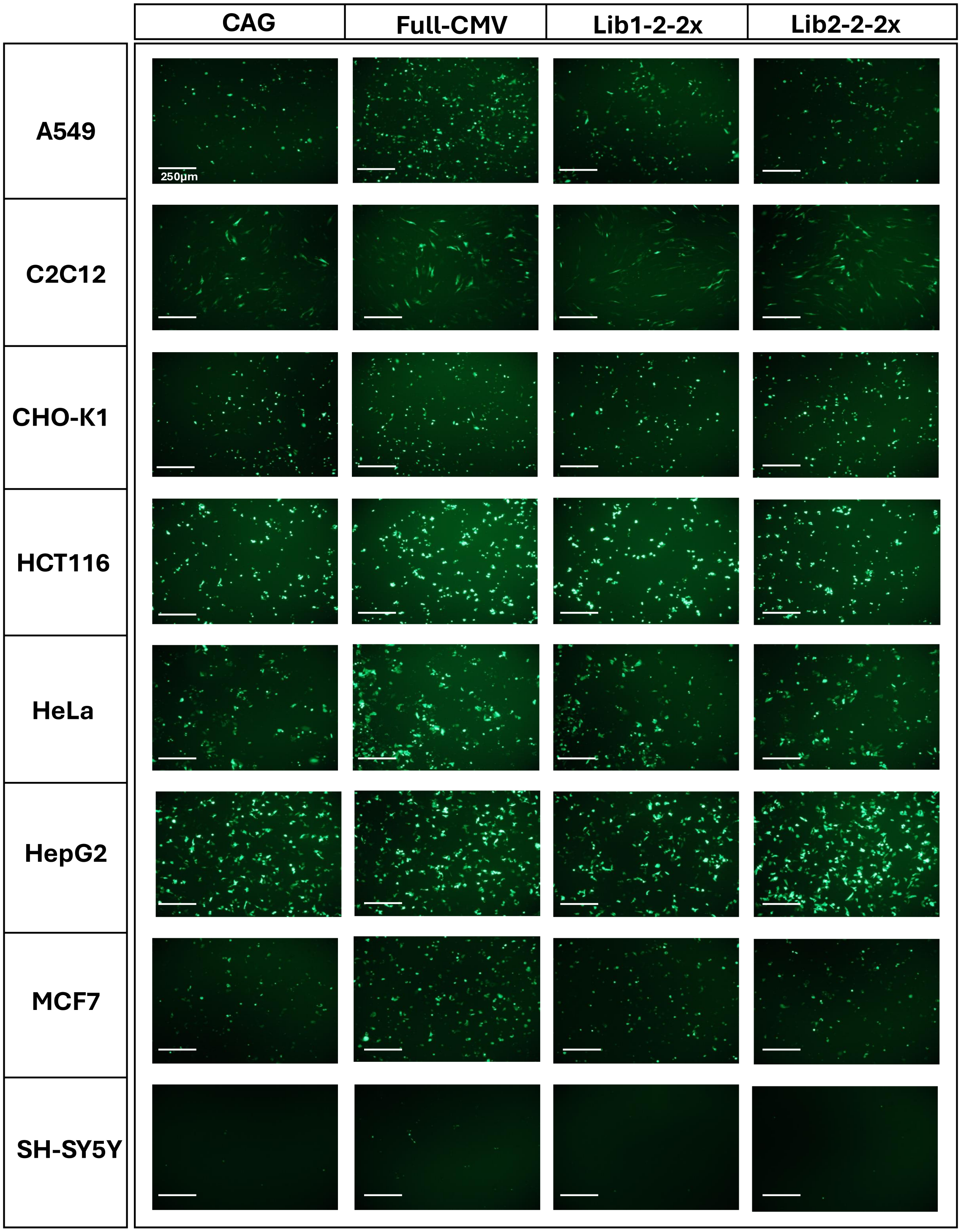


**Supplementary Figure 9. Promoter expression in additional cell lines.** Plasmids containing GFP and CAG, Full-CMV, or the double enhancer variants of Lib1-2 and Lib2-2 were each transfected onto a variety of mammalian cell lines. Images depicting GFP+ cells were captured 48 hrs post-transfection.

#### Supplementary Figure 10

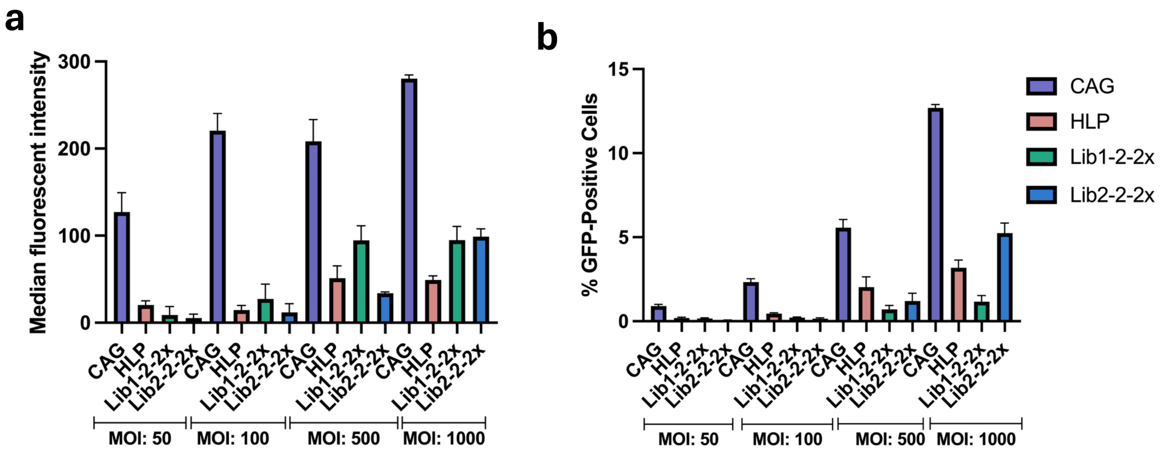


**Supplementary Figure 10. MOI optimization for promoter transduction in liver cells.** To prevent biased dominance of CAG expression due to double- and triple-virion infected cells, HepG2 cells were transduced with AAV2-packaged CAG, HLP, Lib1-2-2x, and Lib2-2-2x at a range of MOIs and fit to a Poisson model. Promoter strength (a) and infectivity rates (b) were assessed by flow cytometry 72 hrs post-transduction. According to the model, only 0.45% of cells are infected with more than one virion at an infectivity rate of $\lambda$=0.1; thus an MOI of 1000 was chosen for subsequent studies. Interestingly, promoter strength and infection efficiency was variable based on MOI, and ELiPS promoters tended to outperform HLP at higher MOIs. Bars represent mean +/- SEM of per-replicate medians.
