## Supplementary Tables for "Expression-linked promoter selection (ELiPS) engineers short, strong ubiquitous promoters for gene therapy applications"

**Supplementary Table 1: Identity and description of transcription factors, binding sites, and barcodes used in ELiPS libraries 1 and 2**

| Library 1 | TF Name | Ensembl | Gene Description | TFBS Sequence | Barcode | Orientation | Top oligo BbsI | Bottom oligo BbsI | Top oligo BsaI | Bottom oligo BsaI |
|---|---|---|---|---|---|---|---|---|---|---|
| | ATF4 | ENSG00000128272 | Activating transcription factor 4 | GGATGATGCAAT | GTCT | Forward | caaaGGATGATGCAATcaaagaGTCTTCaaaGAAGACacactcGTCT | gagtAGACgagtgtGTCTTCtttGAAGACtctttgATTGCATCATCC | caaaGGATGATGCAATcaaagGAGACCaaaGGTCTCcactcGTCT | gagtAGACgagtgGAGACCtttGGTCTCctttgATTGCATCATCC |
| | FOS | ENSG00000170345 | Fos proto-oncogene, AP-1 transcription factor subunit | TGTGACTCAT | CTAC | Forward | caaaTGTGACTCATcaaagaGTCTTCaaaGAAGACacactcCTAC | gagtGTAGgagtgtGTCTTCtttGAAGACtctttgATGAGTCACA | caaaTGTGACTCATcaaagGAGACCaaaGGTCTCcactcCTAC | gagtGTAGgagtgGAGACCtttGGTCTCctttgATGAGTCACA |
| | JUN | ENSG00000177606 | Jun proto-oncogene, AP-1 transcription factor subunit | AAGATGATGTCAT | AGAA | Forward | caaaAAGATGATGTCATcaaagaGTCTTCaaaGAAGACacactcAGAA | gagtTTCTgagtgtGTCTTCtttGAAGACtctttgATGACATCATCTT | caaaAAGATGATGTCATcaaagGAGACCaaaGGTCTCcactcAGAA | gagtTTCTgagtgGAGACCtttGGTCTCctttgATGACATCATCTT |
| | CEBPB | ENSG00000172216 | CCAAT/enhancer binding protein beta | TATTGCACAAT | CCCA | Forward | caaaTATTGCACAATcaaagaGTCTTCaaaGAAGACacactcCCCA | gagtTGGGgagtgtGTCTTCtttGAAGACtctttgATTGTGCAATA | caaaTATTGCACAATcaaagGAGACCaaaGGTCTCcactcCCCA | gagtTGGGgagtgGAGACCtttGGTCTCctttgATTGTGCAATA |
| | NFYA | ENSG00000001167 | Nuclear transcription factor Y subunit alpha | CTCAGCCAATCAGCG | TAAG | Forward | caaaCTCAGCCAATCAGCGcaaagaGTCTTCaaaGAAGACacactcTAAG | gagtCTTAgagtgtGTCTTCtttGAAGACtctttgCGCTGATTGGCTGAG | caaaCTCAGCCAATCAGCGcaaagGAGACCaaaGGTCTCcactcTAAG | gagtCTTAgagtgGAGACCtttGGTCTCctttgCGCTGATTGGCTGAG |
| | EGR1 | ENSG00000120738 | Early growth response 1 | CCCCCGCCCCCGCC | TCGG | Forward | caaaCCCCCGCCCCCGCCcaaagaGTCTTCaaaGAAGACacactcTCGG | gagtCCGAgagtgtGTCTTCtttGAAGACtctttgGGCGGGGGCGGGGG | caaaCCCCCGCCCCCGCCcaaagGAGACCaaaGGTCTCcactcTCGG | gagtCCGAgagtgGAGACCtttGGTCTCctttgGGCGGGGGCGGGGG |
| | NFE2L2 | ENSG00000116044 | Nuclear factor, erythroid 2 like 2 | ATGACTCAGCA | ACCC | Forward | caaaATGACTCAGCAcaaagaGTCTTCaaaGAAGACacactcACCC | gagtGGGTgagtgtGTCTTCtttGAAGACtctttgTGCTGAGTCAT | caaaATGACTCAGCAcaaagGAGACCaaaGGTCTCcactcACCC | gagtGGGTgagtgGAGACCtttGGTCTCctttgTGCTGAGTCAT |
| | NR1H2 | ENSG00000131408 | Nuclear receptor subfamily 1 group H member 2 | AAAGGTCAAAGGTCAAC | CCAA | Forward | caaaAAAGGTCAAAGGTCAACcaaagaGTCTTCaaaGAAGACacactcCCAA | gagtTTGGgagtgtGTCTTCtttGAAGACtctttgGTTGACCTTTGACCTTT | caaaAAAGGTCAAAGGTCAACcaaagGAGACCaaaGGTCTCcactcCCAA | gagtTTGGgagtgGAGACCtttGGTCTCctttgGTTGACCTTTGACCTTT |
| | CREB1 | ENSG00000118260 | CAMP responsive element binding protein 1 | TGACGTCA | TCGT | Forward | caaaTGACGTCAcaaagaGTCTTCaaaGAAGACacactcTCGT | gagtACGAgagtgtGTCTTCtttGAAGACtctttgTGACGTCA | caaaTGACGTCAcaaagGAGACCaaaGGTCTCcactcTCGT | gagtACGAgagtgGAGACCtttGGTCTCctttgTGACGTCA |
| | KLF6 | ENSG00000067082 | Kruppel like factor 6 | GCCACGCCCA | AAGG | Forward | caaaGCCACGCCCAcaaagaGTCTTCaaaGAAGACacactcAAGG | gagtCCTTgagtgtGTCTTCtttGAAGACtctttgTGGGCGTGGC | caaaGCCACGCCCAcaaagGAGACCaaaGGTCTCcactcAAGG | gagtCCTTgagtgGAGACCtttGGTCTCctttgTGGGCGTGGC |
| | ETS2 | ENSG00000157557 | ETS proto-oncogene 2, transcription factor | GACCGGAAGT | GGAT | Forward | caaaGACCGGAAGTcaaagaGTCTTCaaaGAAGACacactcGGAT | gagtATCCgagtgtGTCTTCtttGAAGACtctttgACTTCCGGTC | caaaGACCGGAAGTcaaagGAGACCaaaGGTCTCcactcGGAT | gagtATCCgagtgGAGACCtttGGTCTCctttgACTTCCGGTC |
| | ELF1 | ENSG00000120690 | E74 like ETS transcription factor 1 | GAACCAGGAAGTG | CCTG | Forward | caaaGAACCAGGAAGTGcaaagaGTCTTCaaaGAAGACacactcCCTG | gagtCAGGgagtgtGTCTTCtttGAAGACtctttgCACTTCCTGGTTC | caaaGAACCAGGAAGTGcaaagGAGACCaaaGGTCTCcactcCCTG | gagtCAGGgagtgGAGACCtttGGTCTCctttgCACTTCCTGGTTC |
| | TCF12 | ENSG00000140262 | Transcription factor 12 | CACGTGC | CACG | Forward | caaaCACGTGCcaaagaGTCTTCaaaGAAGACacactcCACG | gagtCGTGgagtgtGTCTTCtttGAAGACtctttgGCACGTG | caaaCACGTGCcaaagGAGACCaaaGGTCTCcactcCACG | gagtCGTGgagtgGAGACCtttGGTCTCctttgGCACGTG |
| | SP1 | ENSG00000185591 | Sp1 transcription factor | GGGGCGGGGT | GTAT | Forward | caaaGGGGCGGGGTcaaagaGTCTTCaaaGAAGACacactcGTAT | gagtATACgagtgtGTCTTCtttGAAGACtctttgACCCCGCCCC | caaaGGGGCGGGGTcaaagGAGACCaaaGGTCTCcactcGTAT | gagtATACgagtgGAGACCtttGGTCTCctttgACCCCGCCCC |
| | POU2F1 | ENSG00000143190 | POU class 2 homeobox 1 | AATATGCAAATAA | GAGA | Forward | caaaAATATGCAAATAAcaaagaGTCTTCaaaGAAGACacactcGAGA | gagtTCTCgagtgtGTCTTCtttGAAGACtctttgTTATTTGCATATT | caaaAATATGCAAATAAcaaagGAGACCaaaGGTCTCcactcGAGA | gagtTCTCgagtgGAGACCtttGGTCTCctttgTTATTTGCATATT |
| | ATF4_rev | ENSG00000128272 | Activating transcription factor 4 | ATTGCATCATCC | AGCC | Reverse | caaaATTGCATCATCCcaaagaGTCTTCaaaGAAGACacactcAGCC | gagtGGCTgagtgtGTCTTCtttGAAGACtctttgGGATGATGCAAT | caaaATTGCATCATCCcaaagGAGACCaaaGGTCTCcactcAGCC | gagtGGCTgagtgGAGACCtttGGTCTCctttgGGATGATGCAAT |
| | FOS_rev | ENSG00000170345 | Fos proto-oncogene, AP-1 transcription factor subunit | ATGAGTCACA | TGTG | Reverse | caaaATGAGTCACAcaaagaGTCTTCaaaGAAGACacactcTGTG | gagtCACAgagtgtGTCTTCtttGAAGACtctttgTGTGACTCAT | caaaATGAGTCACAcaaagGAGACCaaaGGTCTCcactcTGTG | gagtCACAgagtgGAGACCtttGGTCTCctttgTGTGACTCAT |
| | JUN_rev | ENSG00000177606 | Jun proto-oncogene, AP-1 transcription factor subunit | ATGACATCATCTT | TTAT | Reverse | caaaATGACATCATCTTcaaagaGTCTTCaaaGAAGACacactcTTAT | gagtATAAgagtgtGTCTTCtttGAAGACtctttgAAGATGATGTCAT | caaaATGACATCATCTTcaaagGAGACCaaaGGTCTCcactcTTAT | gagtATAAgagtgGAGACCtttGGTCTCctttgAAGATGATGTCAT |
| | CEBPB_rev | ENSG00000172216 | CCAAT/enhancer binding protein beta | ATTGTGCAATA | TACG | Reverse | caaaATTGTGCAATAcaaagaGTCTTCaaaGAAGACacactcTACG | gagtCGTAgagtgtGTCTTCtttGAAGACtctttgTATTGCACAAT | caaaATTGTGCAATAcaaagGAGACCaaaGGTCTCcactcTACG | gagtCGTAgagtgGAGACCtttGGTCTCctttgTATTGCACAAT |
| | NFYA_rev | ENSG00000001167 | Nuclear transcription factor Y subunit alpha | CGCTGATTGGCTGAG | CAGA | Reverse | caaaCGCTGATTGGCTGAGcaaagaGTCTTCaaaGAAGACacactcCAGA | gagtTCTGgagtgtGTCTTCtttGAAGACtctttgCTCAGCCAATCAGCG | caaaCGCTGATTGGCTGAGcaaagGAGACCaaaGGTCTCcactcCAGA | gagtTCTGgagtgGAGACCtttGGTCTCctttgCTCAGCCAATCAGCG |
| | EGR1_rev | ENSG00000120738 | Early growth response 1 | GGCGGGGGCGGGGG | CGAA | Reverse | caaaGGCGGGGGCGGGGGcaaagaGTCTTCaaaGAAGACacactcCGAA | gagtTTCGgagtgtGTCTTCtttGAAGACtctttgCCCCCGCCCCCGCC | caaaGGCGGGGGCGGGGGcaaagGAGACCaaaGGTCTCcactcCGAA | gagtTTCGgagtgGAGACCtttGGTCTCctttgCCCCCGCCCCCGCC |
| | NFE2L2_rev | ENSG00000116044 | Nuclear factor, erythroid 2 like 2 | TGCTGAGTCAT | ATTC | Reverse | caaaTGCTGAGTCATcaaagaGTCTTCaaaGAAGACacactcATTC | gagtGAATgagtgtGTCTTCtttGAAGACtctttgATGACTCAGCA | caaaTGCTGAGTCATcaaagGAGACCaaaGGTCTCcactcATTC | gagtGAATgagtgGAGACCtttGGTCTCctttgATGACTCAGCA |
| | NR1H2_rev | ENSG00000131408 | Nuclear receptor subfamily 1 group H member 2 | GTTGACCTTTGACCTTT | ACAA | Reverse | caaaGTTGACCTTTGACCTTTcaaagaGTCTTCaaaGAAGACacactcACAA | gagtTTGTgagtgtGTCTTCtttGAAGACtctttgAAAGGTCAAAGGTCAAC | caaaGTTGACCTTTGACCTTTcaaagGAGACCaaaGGTCTCcactcACAA | gagtTTGTgagtgGAGACCtttGGTCTCctttgAAAGGTCAAAGGTCAAC |
| | CREB1_rev | ENSG00000118260 | CAMP responsive element binding protein 1 | TGACGTCA | GTCC | Reverse | caaaTGACGTCAcaaagaGTCTTCaaaGAAGACacactcGTCC | gagtGGACgagtgtGTCTTCtttGAAGACtctttgTGACGTCA | caaaTGACGTCAcaaagGAGACCaaaGGTCTCcactcGTCC | gagtGGACgagtgGAGACCtttGGTCTCctttgTGACGTCA |
| | KLF6_rev | ENSG00000067082 | Kruppel like factor 6 | TGGGCGTGGC | AGGC | Reverse | caaaTGGGCGTGGCcaaagaGTCTTCaaaGAAGACacactcAGGC | gagtGCCTgagtgtGTCTTCtttGAAGACtctttgGCCACGCCCA | caaaTGGGCGTGGCcaaagGAGACCaaaGGTCTCcactcAGGC | gagtGCCTgagtgGAGACCtttGGTCTCctttgGCCACGCCCA |
| | ETS2_rev | ENSG00000157557 | ETS proto-oncogene 2, transcription factor | ACTTCCGGTC | TCCT | Reverse | caaaACTTCCGGTCcaaagaGTCTTCaaaGAAGACacactcTCCT | gagtAGGAgagtgtGTCTTCtttGAAGACtctttgGACCGGAAGT | caaaACTTCCGGTCcaaagGAGACCaaaGGTCTCcactcTCCT | gagtAGGAgagtgGAGACCtttGGTCTCctttgGACCGGAAGT |
| | ELF1_rev | ENSG00000120690 | E74 like ETS transcription factor 1 | CACTTCCTGGTTC | CCCT | Reverse | caaaCACTTCCTGGTTCcaaagaGTCTTCaaaGAAGACacactcCCCT | gagtAGGGgagtgtGTCTTCtttGAAGACtctttgGAACCAGGAAGTG | caaaCACTTCCTGGTTCcaaagGAGACCaaaGGTCTCcactcCCCT | gagtAGGGgagtgGAGACCtttGGTCTCctttgGAACCAGGAAGTG |
| | TCF12_rev | ENSG00000140262 | Transcription factor 12 | GCACGTG | GATA | Reverse | caaaGCACGTGcaaagaGTCTTCaaaGAAGACacactcGATA | gagtTATCgagtgtGTCTTCtttGAAGACtctttgCACGTGC | caaaGCACGTGcaaagGAGACCaaaGGTCTCcactcGATA | gagtTATCgagtgGAGACCtttGGTCTCctttgCACGTGC |
| | SP1_rev | ENSG00000185591 | Sp1 transcription factor | ACCCCGCCCC | GTTG | Reverse | caaaACCCCGCCCCcaaagaGTCTTCaaaGAAGACacactcGTTG | gagtCAACgagtgtGTCTTCtttGAAGACtctttgGGGGCGGGGT | caaaACCCCGCCCCcaaagGAGACCaaaGGTCTCcactcGTTG | gagtCAACgagtgGAGACCtttGGTCTCctttgGGGGCGGGGT |
| | POU2F1_rev | ENSG00000143190 | POU class 2 homeobox 1 | TTATTTGCATATT | CGGA | Reverse | caaaTTATTTGCATATTcaaagaGTCTTCaaaGAAGACacactcCGGA | gagtTCCGgagtgtGTCTTCtttGAAGACtctttgAATATGCAAATAA | caaaTTATTTGCATATTcaaagGAGACCaaaGGTCTCcactcCGGA | gagtTCCGgagtgGAGACCtttGGTCTCctttgAATATGCAAATAA |

| Library 2 | TF Name | Ensembl | Gene Description | TFBS Sequence | Barcode | Orientation | Top oligo BbsI | Bottom oligo BbsI | Top oligo BsaI | Bottom oligo BsaI |
|---|---|---|---|---|---|---|---|---|---|---|
| | HNF1A | ENSG00000135100 | HNF1 homeobox A | AGTTAATGATTAACT | TGCG | Forward | caaaAGTTAATGATTAACTcaaagaGTCTTCaaaGAAGACacactcTGCG | gagtCGCAgagtgtGTCTTCtttGAAGACtctttgAGTTAATCATTAACT | caaaAGTTAATGATTAACTcaaagGAGACCaaaGGTCTCcactcTGCG | gagtCGCAgagtgGAGACCtttGGTCTCctttgAGTTAATCATTAACT |
| | HNF1B | ENSG00000275410 | HNF1 homeobox B | GTTAATGATTAAC | TGAA | Forward | caaaGTTAATGATTAACcaaagaGTCTTCaaaGAAGACacactcTGAA | gagtTTCAgagtgtGTCTTCtttGAAGACtctttgGTTAATCATTAAC | caaaGTTAATGATTAACcaaagGAGACCaaaGGTCTCcactcTGAA | gagtTTCAgagtgGAGACCtttGGTCTCctttgGTTAATCATTAAC |
| | HNF4A | ENSG00000101076 | hepatocyte nuclear factor 4, alpha | CTGGACTTTGGCCTC | GGCG | Forward | caaaCTGGACTTTGGCCTCcaaagaGTCTTCaaaGAAGACacactcGGCG | gagtCGCCgagtgtGTCTTCtttGAAGACtctttgGAGGCCAAAGTCCAG | caaaCTGGACTTTGGCCTCcaaagGAGACCaaaGGTCTCcactcGGCG | gagtCGCCgagtgGAGACCtttGGTCTCctttgGAGGCCAAAGTCCAG |
| | NR2F1 | ENSG00000175745 | nuclear receptor subfamily 2, group F, member 1 | CAAAGGTCAAGGA | ACGG | Forward | caaaCAAAGGTCAAGGAcaaagaGTCTTCaaaGAAGACacactcACGG | gagtCCGTgagtgtGTCTTCtttGAAGACtctttgTCCTTGACCTTTG | caaaCAAAGGTCAAGGAcaaagGAGACCaaaGGTCTCcactcACGG | gagtCCGTgagtgGAGACCtttGGTCTCctttgTCCTTGACCTTTG |
| | TBP | ENSG00000112592 | TATA box binding protein | GTATAAAAGGCGGGG | GCGC | Forward | caaaGTATAAAAGGCGGGGcaaagaGTCTTCaaaGAAGACacactcGCGC | gagtGCGCgagtgtGTCTTCtttGAAGACtctttgCCCCGCCTTTTATAC | caaaGTATAAAAGGCGGGGcaaagGAGACCaaaGGTCTCcactcGCGC | gagtGCGCgagtgGAGACCtttGGTCTCctttgCCCCGCCTTTTATAC |
| | FOXA2 | ENSG00000125798 | forkhead box A2 | TGTTTACTTAGG | CGAC | Forward | caaaTGTTTACTTAGGcaaagaGTCTTCaaaGAAGACacactcCGAC | gagtGTCGgagtgtGTCTTCtttGAAGACtctttgCCTAAGTAAACA | caaaTGTTTACTTAGGcaaagGAGACCaaaGGTCTCcactcCGAC | gagtGTCGgagtgGAGACCtttGGTCTCctttgCCTAAGTAAACA |
| | FOXD1 | ENSG00000251493 | forkhead box D1 | GTAAACAT | TAAA | Forward | caaaGTAAACATcaaagaGTCTTCaaaGAAGACacactcTAAA | gagtTTTAgagtgtGTCTTCtttGAAGACtctttgATGTTTAC | caaaGTAAACATcaaagGAGACCaaaGGTCTCcactcTAAA | gagtTTTAgagtgGAGACCtttGGTCTCctttgATGTTTAC |
| | FOXF2 | ENSG00000137273 | forkhead box F2 | CAAACGTAAACAAT | AGTA | Forward | caaaCAAACGTAAACAATcaaagaGTCTTCaaaGAAGACacactcAGTA | gagtTACTgagtgtGTCTTCtttGAAGACtctttgATTGTTTACGTTTG | caaaCAAACGTAAACAATcaaagGAGACCaaaGGTCTCcactcAGTA | gagtTACTgagtgGAGACCtttGGTCTCctttgATTGTTTACGTTTG |
| | FOXA1 | ENSG00000129514 | forkhead box A1 | TCCATGTTTACTTTG | GTTC | Forward | caaaTCCATGTTTACTTTGcaaagaGTCTTCaaaGAAGACacactcGTTC | gagtGAACgagtgtGTCTTCtttGAAGACtctttgCAAAGTAAACATGGA | caaaTCCATGTTTACTTTGcaaagGAGACCaaaGGTCTCcactcGTTC | gagtGAACgagtgGAGACCtttGGTCTCctttgCAAAGTAAACATGGA |
| | PLAG1 | ENSG00000181690 | pleiomorphic adenoma gene 1 | GGGGCCCAAGGGGG | TCTA | Forward | caaaGGGGCCCAAGGGGGcaaagaGTCTTCaaaGAAGACacactcTCTA | gagtTAGAgagtgtGTCTTCtttGAAGACtctttgCCCCCTTGGGCCCC | caaaGGGGCCCAAGGGGGcaaagGAGACCaaaGGTCTCcactcTCTA | gagtTAGAgagtgGAGACCtttGGTCTCctttgCCCCCTTGGGCCCC |
| | GABPA | ENSG00000154727 | GA binding protein transcription factor, alpha subunit 60kDa | CCGGAAGTGGC | CTAA | Forward | caaaCCGGAAGTGGCcaaagaGTCTTCaaaGAAGACacactcCTAA | gagtTTAGgagtgtGTCTTCtttGAAGACtctttgGCCACTTCCGG | caaaCCGGAAGTGGCcaaagGAGACCaaaGGTCTCcactcCTAA | gagtTTAGgagtgGAGACCtttGGTCTCctttgGCCACTTCCGG |
| | MYB | ENSG00000118513 | v-myb myeloblastosis viral oncogene homolog (avian) | ACCAACTGTC | CGAG | Forward | caaaACCAACTGTCcaaagaGTCTTCaaaGAAGACacactcCGAG | gagtCTCGgagtgtGTCTTCtttGAAGACtctttgGACAGTTGGT | caaaACCAACTGTCcaaagGAGACCaaaGGTCTCcactcCGAG | gagtCTCGgagtgGAGACCtttGGTCTCctttgGACAGTTGGT |
| | NFE2L2 | ENSG00000116044 | nuclear factor (erythroid-derived 2)-like 2 | ATGACTCAGCA | TGGC | Forward | caaaATGACTCAGCAcaaagaGTCTTCaaaGAAGACacactcTGGC | gagtGCCAgagtgtGTCTTCtttGAAGACtctttgTGCTGAGTCAT | caaaATGACTCAGCAcaaagGAGACCaaaGGTCTCcactcTGGC | gagtGCCAgagtgGAGACCtttGGTCTCctttgTGCTGAGTCAT |
| | CREB1 | ENSG00000118260 | cAMP responsive element binding protein 1 | GGTGACGTCACC | GTTT | Forward | caaaGGTGACGTCACCcaaagaGTCTTCaaaGAAGACacactcGTTT | gagtAAACgagtgtGTCTTCtttGAAGACtctttgGGTGACGTCACC | caaaGGTGACGTCACCcaaagGAGACCaaaGGTCTCcactcGTTT | gagtAAACgagtgGAGACCtttGGTCTCctttgGGTGACGTCACC |
| | EGR1 | ENSG00000120738 | Early Growth Response 1 | TACGCCCACGCATT | TGGT | Forward | caaaTACGCCCACGCATTcaaagaGTCTTCaaaGAAGACacactcTGGT | gagtACCAgagtgtGTCTTCtttGAAGACtctttgAATGCGTGGGCGTA | caaaTACGCCCACGCATTcaaagGAGACCaaaGGTCTCcactcTGGT | gagtACCAgagtgGAGACCtttGGTCTCctttgAATGCGTGGGCGTA |
| | E2F1 | ENSG00000101412 | E2F1 | TTTGGCGCCAAA | AGAT | Forward | caaaTTTGGCGCCAAAcaaagaGTCTTCaaaGAAGACacactcAGAT | gagtATCTgagtgtGTCTTCtttGAAGACtctttgTTTGGCGCCAAA | caaaTTTGGCGCCAAAcaaagGAGACCaaaGGTCTCcactcAGAT | gagtATCTgagtgGAGACCtttGGTCTCctttgTTTGGCGCCAAA |
| | NRF1 | ENSG00000106459 | Nuclear respiratory factor 1 | TGCGCACGCGCA | GGGA | Forward | caaaTGCGCACGCGCAcaaagaGTCTTCaaaGAAGACacactcGGGA | gagtTCCCgagtgtGTCTTCtttGAAGACtctttgTGCGCGTGCGCA | caaaTGCGCACGCGCAcaaagGAGACCaaaGGTCTCcactcGGGA | gagtTCCCgagtgGAGACCtttGGTCTCctttgTGCGCGTGCGCA |
| | HNF1A_rev | ENSG00000135100 | HNF1 homeobox A | AGTTAATCATTAACT | CTAG | Reverse | caaaAGTTAATCATTAACTcaaagaGTCTTCaaaGAAGACacactcCTAG | gagtCTAGgagtgtGTCTTCtttGAAGACtctttgAGTTAATGATTAACT | caaaAGTTAATCATTAACTcaaagGAGACCaaaGGTCTCcactcCTAG | gagtCTAGgagtgGAGACCtttGGTCTCctttgAGTTAATGATTAACT |
| | HNF1B_rev | ENSG00000275410 | HNF1 homeobox B | GTTAATCATTAAC | TACA | Reverse | caaaGTTAATCATTAACcaaagaGTCTTCaaaGAAGACacactcTACA | gagtTGTAgagtgtGTCTTCtttGAAGACtctttgGTTAATGATTAAC | caaaGTTAATCATTAACcaaagGAGACCaaaGGTCTCcactcTACA | gagtTGTAgagtgGAGACCtttGGTCTCctttgGTTAATGATTAAC |
| | HNF4A_rev | ENSG00000101076 | hepatocyte nuclear factor 4, alpha | GAGGCCAAAGTCCAG | GATT | Reverse | caaaGAGGCCAAAGTCCAGcaaagaGTCTTCaaaGAAGACacactcGATT | gagtAATCgagtgtGTCTTCtttGAAGACtctttgCTGGACTTTGGCCTC | caaaGAGGCCAAAGTCCAGcaaagGAGACCaaaGGTCTCcactcGATT | gagtAATCgagtgGAGACCtttGGTCTCctttgCTGGACTTTGGCCTC |
| | NR2F1_rev | ENSG00000175745 | nuclear receptor subfamily 2, group F, member 1 | TCCTTGACCTTTG | GAAA | Reverse | caaaTCCTTGACCTTTGcaaagaGTCTTCaaaGAAGACacactcGAAA | gagtTTTCgagtgtGTCTTCtttGAAGACtctttgCAAAGGTCAAGGA | caaaTCCTTGACCTTTGcaaagGAGACCaaaGGTCTCcactcGAAA | gagtTTTCgagtgGAGACCtttGGTCTCctttgCAAAGGTCAAGGA |
| | TBP_rev | ENSG00000112592 | TATA box binding protein | CCCCGCCTTTTATAC | AAGA | Reverse | caaaCCCCGCCTTTTATACcaaagaGTCTTCaaaGAAGACacactcAAGA | gagtTCTTgagtgtGTCTTCtttGAAGACtctttgGTATAAAAGGCGGGG | caaaCCCCGCCTTTTATACcaaagGAGACCaaaGGTCTCcactcAAGA | gagtTCTTgagtgGAGACCtttGGTCTCctttgGTATAAAAGGCGGGG |
| | FOXA2_rev | ENSG00000125798 | forkhead box A2 | CCTAAGTAAACA | GTGA | Reverse | caaaCCTAAGTAAACAcaaagaGTCTTCaaaGAAGACacactcGTGA | gagtTCACgagtgtGTCTTCtttGAAGACtctttgTGTTTACTTAGG | caaaCCTAAGTAAACAcaaagGAGACCaaaGGTCTCcactcGTGA | gagtTCACgagtgGAGACCtttGGTCTCctttgTGTTTACTTAGG |
| | FOXD1_rev | ENSG00000251493 | forkhead box D1 | ATGTTTAC | AAGC | Reverse | caaaATGTTTACcaaagaGTCTTCaaaGAAGACacactcAAGC | gagtGCTTgagtgtGTCTTCtttGAAGACtctttgGTAAACAT | caaaATGTTTACcaaagGAGACCaaaGGTCTCcactcAAGC | gagtGCTTgagtgGAGACCtttGGTCTCctttgGTAAACAT |
| | FOXF2_rev | ENSG00000137273 | forkhead box F2 | ATTGTTTACGTTTG | GGTT | Reverse | caaaATTGTTTACGTTTGcaaagaGTCTTCaaaGAAGACacactcGGTT | gagtAACCgagtgtGTCTTCtttGAAGACtctttgCAAACGTAAACAAT | caaaATTGTTTACGTTTGcaaagGAGACCaaaGGTCTCcactcGGTT | gagtAACCgagtgGAGACCtttGGTCTCctttgCAAACGTAAACAAT |
| | FOXA1_rev | ENSG00000129514 | forkhead box A1 | CAAAGTAAACATGGA | GGGT | Reverse | caaaCAAAGTAAACATGGAcaaagaGTCTTCaaaGAAGACacactcGGGT | gagtACCCgagtgtGTCTTCtttGAAGACtctttgTCCATGTTTACTTTG | caaaCAAAGTAAACATGGAcaaagGAGACCaaaGGTCTCcactcGGGT | gagtACCCgagtgGAGACCtttGGTCTCctttgTCCATGTTTACTTTG |
| | PLAG1_rev | ENSG00000181690 | pleiomorphic adenoma gene 1 | CCCCCTTGGGCCCC | TCAA | Reverse | caaaCCCCCTTGGGCCCCcaaagaGTCTTCaaaGAAGACacactcTCAA | gagtTTGAgagtgtGTCTTCtttGAAGACtctttgGGGGCCCAAGGGGG | caaaCCCCCTTGGGCCCCcaaagGAGACCaaaGGTCTCcactcTCAA | gagtTTGAgagtgGAGACCtttGGTCTCctttgGGGGCCCAAGGGGG |
| | GABPA_rev | ENSG00000154727 | GA binding protein transcription factor, alpha subunit 60kDa | GCCACTTCCGG | GGCA | Reverse | caaaGCCACTTCCGGcaaagaGTCTTCaaaGAAGACacactcGGCA | gagtTGCCgagtgtGTCTTCtttGAAGACtctttgCCGGAAGTGGC | caaaGCCACTTCCGGcaaagGAGACCaaaGGTCTCcactcGGCA | gagtTGCCgagtgGAGACCtttGGTCTCctttgCCGGAAGTGGC |
| | MYB_rev | ENSG00000118513 | v-myb myeloblastosis viral oncogene homolog (avian) | GACAGTTGGT | CAAT | Reverse | caaaGACAGTTGGTcaaagaGTCTTCaaaGAAGACacactcCAAT | gagtATTGgagtgtGTCTTCtttGAAGACtctttgACCAACTGTC | caaaGACAGTTGGTcaaagGAGACCaaaGGTCTCcactcCAAT | gagtATTGgagtgGAGACCtttGGTCTCctttgACCAACTGTC |
| | NFE2L2_rev | ENSG00000116044 | nuclear factor (erythroid-derived 2)-like 2 | TGCTGAGTCAT | GACA | Reverse | caaaTGCTGAGTCATcaaagaGTCTTCaaaGAAGACacactcGACA | gagtTGTCgagtgtGTCTTCtttGAAGACtctttgATGACTCAGCA | caaaTGCTGAGTCATcaaagGAGACCaaaGGTCTCcactcGACA | gagtTGTCgagtgGAGACCtttGGTCTCctttgATGACTCAGCA |
| | CREB1_rev | ENSG00000118260 | cAMP responsive element binding protein 1 | GGTGACGTCACC | CTCG | Reverse | caaaGGTGACGTCACCcaaagaGTCTTCaaaGAAGACacactcCTCG | gagtCGAGgagtgtGTCTTCtttGAAGACtctttgGGTGACGTCACC | caaaGGTGACGTCACCcaaagGAGACCaaaGGTCTCcactcCTCG | gagtCGAGgagtgGAGACCtttGGTCTCctttgGGTGACGTCACC |
| | EGR1_rev | ENSG00000120738 | Early Growth Response 1 | AATGCGTGGGCGTA | ATTT | Reverse | caaaAATGCGTGGGCGTAcaaagaGTCTTCaaaGAAGACacactcATTT | gagtAAATgagtgtGTCTTCtttGAAGACtctttgTACGCCCACGCATT | caaaAATGCGTGGGCGTAcaaagGAGACCaaaGGTCTCcactcATTT | gagtAAATgagtgGAGACCtttGGTCTCctttgTACGCCCACGCATT |
| | E2F1_rev | ENSG00000101412 | E2F1 | TTTGGCGCCAAA | CCTT | Reverse | caaaTTTGGCGCCAAAcaaagaGTCTTCaaaGAAGACacactcCCTT | gagtAAGGgagtgtGTCTTCtttGAAGACtctttgTTTGGCGCCAAA | caaaTTTGGCGCCAAAcaaagGAGACCaaaGGTCTCcactcCCTT | gagtAAGGgagtgGAGACCtttGGTCTCctttgTTTGGCGCCAAA |
| | NRF1_rev | ENSG00000106459 | Nuclear respiratory factor 1 | TGCGCGTGCGCA | TTGG | Reverse | caaaTGCGCGTGCGCAcaaagaGTCTTCaaaGAAGACacactcTTGG | gagtCCAAgagtgtGTCTTCtttGAAGACtctttgTGCGCACGCGCA | caaaTGCGCGTGCGCAcaaagGAGACCaaaGGTCTCcactcTTGG | gagtCCAAgagtgGAGACCtttGGTCTCctttgTGCGCACGCGCA |

**Supplementary Table 2: Identity of the top 3 promoters from each library used in studies.**

| Library | Promoter Name | Promoter Size (with SCP2) | BC1 | BC2 | BC3 | BC4 | BC5 | BC6 | BC7 | BC8 | Promoter Sequence (with SCP2 ) | Promoter Sequence with double enhancer, if applicable |
|---|---|---|---|---|---|---|---|---|---|---|---|---|
| 1 | Lib1-1 | 193 | JUN_rev | NFE2L2_rev | EGR1 | KLF6_rev | NFYA | SP1_rev | CEBPB | | ATGACATCATCTTCAAATGCTGAGTCATCAAACCCCCGCCCCCGCCCAAATGGGCGTGGCCAAACTCAGCCAATCAGCGCAAAACCCCGCCCCCAAATATTGCACAATcaaaAGGTCTATATAAGCAGAGCTCGTTTAGTGAACCGTCAGATCGCCTGGAGACGTCGAGCCGAGTGGTTGTGCCTCCATAGAA | |
| 1 | Lib1-2 | 193 | NR1H2_rev | POU2F1 | TCF12_rev | ATF4_rev | FOS_rev | JUN_rev | ATF4_rev | | GTTGACCTTTGACCTTTCAAAAATATGCAAATAACAAAGCACGTGCAAAATTGCATCATCCCAAAATGAGTCACACAAAATGACATCATCTTCAAAATTGCATCATCCcaaaAGGTCTATATAAGCAGAGCTCGTTTAGTGAACCGTCAGATCGCCTGGAGACGTCGAGCCGAGTGGTTGTGCCTCCATAGAA | GTTGACCTTTGACCTTTCAAAAATATGCAAATAACAAAGCACGTGCAAAATTGCATCATCCCAAAATGAGTCACACAAAATGACATCATCTTCAAAATTGCATCATCCGAACGTTGACCTTTGACCTTTCAAAAATATGCAAATAACAAAGCACGTGCAAAATTGCATCATCCCAAAATGAGTCACACAAAATGACATCATCTTCAAAATTGCATCATCCcaaaAGGTCTATATAAGCAGAGCTCGTTTAGTGAACCGTCAGATCGCCTGGAGACGTCGAGCCGAGTGGTTGTGCCTCCATAGAA |
| 1 | Lib1-3 | 201 | NFE2L2 | CREB1_rev | CEBPB | FOS_rev | SP1_rev | ATF4_rev | JUN_rev | POU2F1_rev | ATGACTCAGCACAAATGACGTCACAAATATTGCACAATCAAAATGAGTCACACAAAACCCCGCCCCCAAAATTGCATCATCCCAAAATGACATCATCTTCAAATTATTTGCATATTcaaaAGGTCTATATAAGCAGAGCTCGTTTAGTGAACCGTCAGATCGCCTGGAGACGTCGAGCCGAGTGGTTGTGCCTCCATAGAA | |
| 2 | Lib2-1 | 213 | FOXA1_rev | FOXF2_rev | FOXD1_rev | NR2F1_rev | GABPA | EGR1 | GABPA | | CAAAGTAAACATGGACAAAATTGTTTACGTTTGCAAAATGTTTACCAAATCCTTGACCTTTGCAAACCGGAAGTGGCCAAATACGCCCACGCATTCAAATACGCCCACGCATTCAAACCGGAAGTGGCcaaaAGGTCTATATAAGCAGAGCTCGTTTAGTGAACCGTCAGATCGCCTGGAGACGTCGAGCCGAGTGGTTGTGCCTCCATAGAA | |
| 2 | Lib2-2 | 218 | EGR1 | HNF1A_rev | NR2F1_rev | E2F1_rev | CREB1_rev | NFE2L2_rev | FOXF2 | TBP | TACGCCCACGCATTCAAAAGTTAATCATTAACTCAAATGCGCGTGCGCACAAATTTGGCGCCAAACAAAGGTGACGTCACCCAAATGCTGAGTCATCAAACAAACGTAAACAATCAAAGTATAAAAGGCGGGGcaaaAGGTCTATATAAGCAGAGCTCGTTTAGTGAACCGTCAGATCGCCTGGAGACGTCGAGCCGAGTGGTTGTGCCTCCATAGAA | TACGCCCACGCATTCAAAAGTTAATCATTAACTCAAATGCGCGTGCGCACAAATTTGGCGCCAAACAAAGGTGACGTCACCCAAATGCTGAGTCATCAAACAAACGTAAACAATCAAAGTATAAAAGGCGGGGGAACTACGCCCACGCATTCAAAAGTTAATCATTAACTCAAATGCGCGTGCGCACAAATTTGGCGCCAAACAAAGGTGACGTCACCCAAATGCTGAGTCATCAAACAAACGTAAACAATCAAAGTATAAAAGGCGGGGcaaaAGGTCTATATAAGCAGAGCTCGTTTAGTGAACCGTCAGATCGCCTGGAGACGTCGAGCCGAGTGGTTGTGCCTCCATAGAA |
| 2 | Lib2-3 | 175 | NR2F1_rev | NFE2L2 | NFE2L2 | NR2F1_rev | NFE2L2 | NFE2L2_rev | | | TCCTTGACCTTTGCAAAATGACTCAGCACAAAATGACTCAGCACAAATCCTTGACCTTTGCAAAATGACTCAGCACAAATGCTGAGTCATcaaaAGGTCTATATAAGCAGAGCTCGTTTAGTGAACCGTCAGATCGCCTGGAGACGTCGAGCCGAGTGGTTGTGCCTCCATAGAA | |

**Supplementary Table 3: Sequences of constitutive promoters used in studies**

| Promoter Name | Length | Sequence |
|---|---|---|
| CAG | 1664 | actagttattaatagtaatcaattacggggtcattagttcatagcccatatatggagttccgcgttacataacttacggtaaatggcccgcctggctgaccgcccaacgacccccgcccattgacgtcaataatgacgtatgttcccatagtaacgccaatagggactttccattgacgtcaatgggtggagtatttacggtaaactgcccacttggcagtacatcaagtgtatcatatgccaagtacgccccctattgacgtcaatgacggtaaatggcccgcctggcattatgcccagtacatgaccttatgggactttcctacttggcagtacatctacgtattagtcatcgctattaccatggtcgaggtgagccccacgttctgcttcactctccccatctcccccccctccccacccccaattttgtatttatttattttttaattattttgtgcagcgatgggggcgggggggggggggggggcgcgcgccaggcggggcggggcggggcgaggggcggggcggggcgaggcggagaggtgcggcggcagccaatcagagcggcgcgctccgaaagtttccttttatggcgaggcggcggcggcggcggccctataaaaagcgaagcgcgcggcgggcggggagtcgctgcgacgctgccttcgccccgtgccccgctccgccgccgcctcgcgccgcccgccccggctctgactgaccgcgttactcccacaggtgagcgggcgggacggcccttctcctccgggctgtaattagcgcttggtttaatgacggcttgtttcttttctgtggctgcgtgaaagccttgaggggctccgggagggccctttgtgcggggggagcggctcggggggtgcgtgcgtgtgtgtgtgcgtggggagcgccgcgtgcggctccgcgctgcccggcggctgtgagcgctgcgggcgcggcgcggggctttgtgcgctccgcagtgtgcgcgaggggagcgcggccgggggcggtgccccgcggtgcggggggggctgcgaggggaacaaaggctgcgtgcggggtgtgtgcgtgggggggtgagcagggggtgtgggcgcgtcggtcgggctgcaaccccccctgcacccccctccccgagttgctgagcacggcccggcttcgggtgcggggctccgtacggggcgtggcgcggggctcgccgtgccgggcggggggtggcggcaggtgggggtgccgggcggggcggggccgcctcgggccggggagggctcgggggaggggcgcggcggcccccggagcgccggcggctgtcgaggcgcggcgagccgcagccattgccttttatggtaatcgtgcgagagggcgcagggacttcctttgtcccaaatctgtgcggagccgaaatctgggaggcgccgccgcaccccctctagcgggcgcggggcgaagcggtgcggcgccggcaggaaggaaatgggcggggagggccttcgtgcgtcgccgcgccgccgtccccttctccctctccagcctcggggctgtccgcggggggacggctgccttcgggggggacggggcagggcggggttcggcttctggcgtgtgaccggcggctctagagcctctgctaaccatgttcatgccttcttctttttcctacag |
| Mini-CMV | 173 | ACTCACGGGGATTTCCAAGTCTCCACCCCATTGACGTCAATGGGAGTTTGTTTTGGCACCAAAATCAACGGGACTTTCCAAAATGTCGTAATAACCCCGCCCCGTTGACGCAAATGGGCGGTAGGCGTGTACGGTGGGAGGTCTATATAAGCAGAGCTCGTTTAGTGAACCGT |
| Full-CMV | 811 | Acattgattattgactagttattaatagtaatcaattacggggtcattagttcatagcccatatatggagttccgcgttacataacttacggtaaatggcccgcctggctgaccgcccaacgacccccgcccattgacgtcaataatgacgtatgttcccatagtaacgccaatagggactttccattgacgtcaatgggtggagtatttacggtaaactgcccacttggcagtacatcaagtgtatcatatgccaagtccgccccctattgacgtcaatgacggtaaatggcccgcctggcattatgcccagtacatgaccttacgggactttcctacttggcagtacatctacgtattagtcatcgctattaccatggtgatgcggttttggcagtacaccaatgggcgtggatagcggtttgactcacggggatttccaagtctccaccccattgacgtcaatgggagtttgttttggcaccaaaatcaacgggactttccaaaatgtcgtaataaccccgccccgttgacgcaaatgggcggtaggcgtgtacggtgggaggtctatataagcagaggtcgtttagtgaaccgtcagatcactagtagctttattgcggtagtttatcacagttaaattgctaacgcagtcagtgctcgactgatcacaggtaagtatcaaggttacaagacaggtttaaggaggccaatagaaactgggcttgtcgagacagagaagattcttgcgtttctgataggcacctattggtcttactgacatccactttgcctttctctccacagggcg |
| HLP | 252 | tgtttgctgcttgcaatgtttgcccattttagggtggacacaggacgctgtggtttctgagccagggggcgactcagatcccagccagtggacttagcccctgtttgctcctccgataactggggtgaccttggttaatattcaccagcagcctcccccgttgcccctctggatccactgcttaaatacggacgaggacagggccctgtctcctcagcttcaggcaccaccactgacctgggacagtgaatc |

**Supplementary Table 4: Introns tested in top promoters**

| Name | Size | Sequence |
|---|---|---|
| Full-CMV Intron | 228 | cgtttagtgaaccgtcagatcactagtagctttattgcggtagtttatcacagttaaattgctaacgcagtcagtgctcgactgatcacaggtaagtatcaaggttacaagacaggtttaaggaggccaatagaaactgggcttgtcgagacagagaagattcttgcgtttctgataggcacctattggtcttactgacatccactttgcctttctctccacagggcg |
| Mini-CMV Intron | 133 | gtaagtatcaaggttacaagacaggtttaaggaggccaatagaaactgggcttgtcgagacagagaagattcttgcgtttctgataggcacctattggtcttactgacatccactttgcctttctctccacag |
| SV40 Intron | 93 | ctctaaggtaaatataaaatttttaagtgtataatgtgttaaactactgattctaattgtttctctcttttagattccaacctttggaactga |

**Supplementary Table 5: Modifications to BDDFVIII gene used in studies**

| Original BDDFVIII sequence | Length | X5 Modifications | Resource |
|---|---|---|---|
| atgcaaatagagctctccacctgcttctttctgtgccttttgcgattctgctttagtgccaccagaagatactacctgggtgcagtggaactgtcatgggactatatgcaaagtgatctcggt<br>gagctgcctgtggacgcaagatttcctcctagagtgccaaaatcttttccattcaacacctcagtcgtgtacaaaaagactctgtttgtagaattcacggatcaccttttcaacatcgcta<br>agccaaggccaccctggatgggtctgctaggtcctaccatccaggctgaggtttatgatacagtggtcattacacttaagaacatggcttcccatcctgtcagtcttcatgctgttggtgtat<br>cctactggaaagcttctgagggagctgaatatgatgatcagaccagtcaaagggagaaagaagatgataaagtcttccctggtggaagccatacatatgtctggcaggtcctgaaagaga<br>atggtccaatggcctctgacccactgtgccttacctactcatatctttctcatgtggacctggtaaaagacttgaattcaggcctcattggagccctactagtatgtagagaagggagtctg<br>gccaaggaaaagacacagaccttgcacaaatttatactactttttgctgtatttgatgaagggaaaagttggcactcagaaacaaagaactccttgatgcaggatagggatgctgcatct<br>gctcgggcctggcctaaaatgcacacagtcaatggttatgtaaacaggtctctgccaggtctgattggatgccacaggaaatcagtctattggcatgtgattggaatgggcaccactcctg<br>aagtgcactcaatattcctcgaaggtcacacatttcttgtgaggaaccatcgccaggcgtccttggaaatctcgccaataactttccttactgctcaaacactcttgatggaccttggac<br>agtttctactgttttgtcatatctcttcccaccaacatgatggcatggaagcttatgtcaaagtagacagctgtccagaggaaccccaactacgaatgaaaaataatgaagaagcggaag<br>actatgatgatgatcttactgattctgaaatggatgtggtcaggtttgatgatgacaactctccttcctttatccaaattcgctcagttgccaagaagcatcctaaaacttgggtacattaca<br>ttgctgctgaagaggaggactgggactatgctcccttagtcctcgcccccgatgacagaagttataaaagtcaatatttgaacaatggccctcagcggattggtaggaagtacaaaaaag<br>tccgatttatggcatacacagatgaaacctttaagactcgtgaagctattcagcatgaatcaggaatcttgggacctttactttatggggaagttggagacacactgttgattatatttaaga<br>atcaagcaagcagaccatataacatctaccctcacggaatcactgatgtccgtcctttgtattcaaggagattaccaaaaggtgtaaaacatttgaaggattttccaattctgccaggag<br>aaatattcaaatataaatggacagtgactgtagaagatgggccaactaaatcagatcctcggtgcctgacccgctattactctagtttcgttaatatggagagagatctagcttcaggact<br>cattggccctctcctcatctgctacaaagaatctgtagatcaaagaggaaaccagataatgtcagacaagaggaatgtcatcctgttttctgtatttgatgagaaccgaagctggtacctc<br>acagagaatatacaacgctttctccccaatccagctggagtgcagcttgaggatccagagttccaagcctccaacatcatgcacagcatcaatggctatgtttttgatagtttgcagttgt<br>cagtttgtttgcatgaggtggcatactggtacattctaagcattggagcacagactgacttcctttctgtcttcttctctggatataccttcaaacacaaaatggtctatgaagacacactca<br>ccctattcccattctcaggagaaactgtcttcatgtcgatggaaaacccaggtctatggattctggggtgccacaactcagactttcggaacagaggcatgaccgccttactgaaggtttc<br>tagttgtgacaagaacactggtgattattacgaggacagttatgaagatatttcagcatacttgctgagtaaaaacaatgccattgaaccaagaagcttctctcaaaacccaccagtctt<br>gaaacgccatcaacgggaaataactcgtactactcttcagtcagatcaagaggaaattgactatgatgataccatatcagttgaaatgaagaaggaagattttgacatttatgatgagga<br>tgaaaatcagagcccccgcagctttcaaaagaaaacacgacactattttattgctgcagtggagaggctctgggattatgggatgagtagctccccacatgttctaagaaacagggctca<br>gagtggcagtgtccctcagttcaagaaagttgttttccaggaatttactgatggctcctttactcagcccttataccgtggagaactaaatgaacatttgggactcctggggccatatataag<br>agcagaagttgaagataatatcatggtaactttcagaaatcaggcctctcgtccctattccttctattctagccttatttcttatgaggaagatcagaggcaaggagcagaacctagaaaa<br>aactttgtcaagcctaatgaaaccaaaacttacttttggaaagtgcaacatcatatggcacccactaaagatgagtttgactgcaaagcctgggcttatttctctgatgttgacctggaaa<br>aagatgtgcactcaggcctgattggaccccttctggtctgccacactaacacactgaaccctgctcatgggagacaagtgacagtacaggaatttgctctgtttttcaccatctttgatga<br>gaccaaaagctggtacttcactgaaaatatggaaagaaactgcagggctccctgcaatatccagatggaagatcccacttttaaagagaattatcgcttccatgcaatcaatggctaca | 4374 | V86I, S108A, K132G, T147M, P152L | Cao, W. *et al.* Molecular Therapy: Methods & Clinical Development (2020). |

| Minimal polyA sequence | Length | Resource |
|---|---|---|
| aataaaagatctttattttcattagatctgtgtgttggttttttgtgtg | 49 | Choi, J.-H. *et al.* Molecular Brain (2014). |
